# Interdigitating coiled-coil tetramers define the helical architecture of GFAP filaments in astrocytes

**DOI:** 10.64898/2026.02.23.707361

**Authors:** Charlie T. Beales, Matthias Eibauer, Yolanda de Pablo, Simon B. Hansen, Ni-Hsuan Lin, Ming-Der Perng, Milos Pekny, Ohad Medalia

## Abstract

GFAP filaments are core elements of the cytoskeleton in astrocytes, shaping cellular processes and influencing central nervous system function and pathology. How these filaments are built at the molecular level has remained unclear. Using cryo-electron microscopy and cryo-electron tomography, we show that GFAP filaments are helical polymers assembled from extended and twisted coiled-coil tetramers that interdigitate to form a complex filament tube. In the filament lumen, the low-complexity head domains aggregate to form a flexible fibre, while the tail domains extend from the surface and facilitate higher-order bundling of the filaments, indicating that both the low-complexity structural regions and the well-ordered coiled-coils of the filament tube contribute to the unique structure of GFAP filaments. Our findings provide molecular insights into the pathogenesis of Alexander Disease, localising multiple deleterious mutations to the critical interlock interaction between successive tetramers emergent in the final filament assembly.

## Introduction

Glial fibrillary acidic protein (GFAP) is the canonical marker of mature astrocytes^1,2^. The most abundant isoform, GFAP-α, self-assembles into intermediate filaments^3–6^ (IFs) that form an extensive cytoskeletal network in astrocytes. GFAP filaments contribute to diverse central nervous system (CNS) functions, including glial motility and proliferation^7–10^, neurogenesis and memory^11–13^, resistance to mechanical and ischemic stress^14–19^, post-traumatic healing^20,21^, regeneration of neuronal axons and synapses after trauma^21–23^, and reorganization of neuronal connections after stroke^24^. The upregulation of GFAP is an important hallmark of astrocyte reactivity in CNS trauma, stroke or neurodegenerative diseases^25^. Mutations in GFAP are responsible for Alexander Disease (AxD), a severe leukodystrophy originating in astrocytes^26^, which is characterised by GFAP aggregation into Rosenthal fibers resulting from filament assembly defects^4,27–30^.

GFAP monomers consist of a highly conserved rod domain flanked by intrinsically disordered N-terminal head and C-terminal tail domains^31–33^. The rod domain is partitioned into the α-helical 1A, 1B, and Coil-2 domains, which are connected by the flexible, non-helical linkers L1 and L12, respectively. GFAP assembly initiates by the formation of α-helical coiled-coil dimers, which interact to form anti-parallel tetramers. These extended tetramers (~65 nm in length) are the fundamental building block for GFAP filament assembly^6,34,35^. Lateral and longitudinal tetramer association produces mature ~10 nm diameter GFAP filaments^36–38^. Highly conserved sequences at both ends of the GFAP rod domain are known to be essential for filament assembly, including the LNDR motif in the 1A domain^27^ and the TYRKLLEGE sequence at the end of Coil-2 domain^39^. Mutations at these regions are associated with severe forms of AxD^27,40^.

Due to the elongated, flexible subunits^41,42^ and substantial polymorphism^37,38,43–45^ determining the structures of assembled IFs has remained a formidable challenge^46,47^. Assembled IFs are not amenable to crystallography, restricting available crystal structures to isolated rod domain fragments^41,48–50^. Only one such crystal structure of the 1B coiled-coil domain has been determined for GFAP^51^. Recently, the first cryo-electron microscopy (cryo-EM) structure of a fully assembled IF was determined^52^. In that study, vimentin (VIM) filaments were shown to form a helical assembly of 40 VIM polypeptides packed into a pentameric cross-section that form a tube-like structure, integrating the low-complexity head domains into the lumen of the filaments.

Here, we used cryo-EM and cryo-electron tomography (cryo-ET) to investigate in situ polymerised GFAP filaments within primary astrocytes. Through power spectrum analysis and helical reconstruction, we resolved their 3D structure. Building on these results, we generated the first molecular model of fully assembled GFAP filaments. This model exhibits a complex helical architecture composed of 32 polypeptides in cross-section, arranged into twisted and interdigitating GFAP tetramers. The low-complexity head domains integrate into the filament lumen as a flexible fiber, while the tail domains extend from the surface, and are engaged in bundling of the filaments within astrocytes. Our model suggests AxD associated mutations spatially cluster at the interlock region^52^. This structural entity emerges from the highly conserved sequences at both ends of the GFAP rod domain as a result of higher-order tetramer interactions in the fully assembled GFAP filaments.

## Results

### Cell-polymerised GFAP filaments are helical polymers

To study GFAP filaments by cryo-EM, we cultured passage 1 primary mouse astrocytes on EM grids. To ensure identification of GFAP filaments, we eliminated VIM filaments by using cells from vimentin null (*Vim^−/−^*) mice^53,54^ **(Fig. 1A)**. To increase the amount of GFAP filaments to the numbers required for cryo-EM analysis, we treated the cells with db-CAMP, inducing a state similar to reactive gliosis^55^. Treatment with 250 µM db-CAMP for 14 days resulted in a striking increase in GFAP immunoreactivity **(Fig. 1B)** and extension of a large number of astrocytic processes **(Fig. 1C)**.

**Fig. 1:**
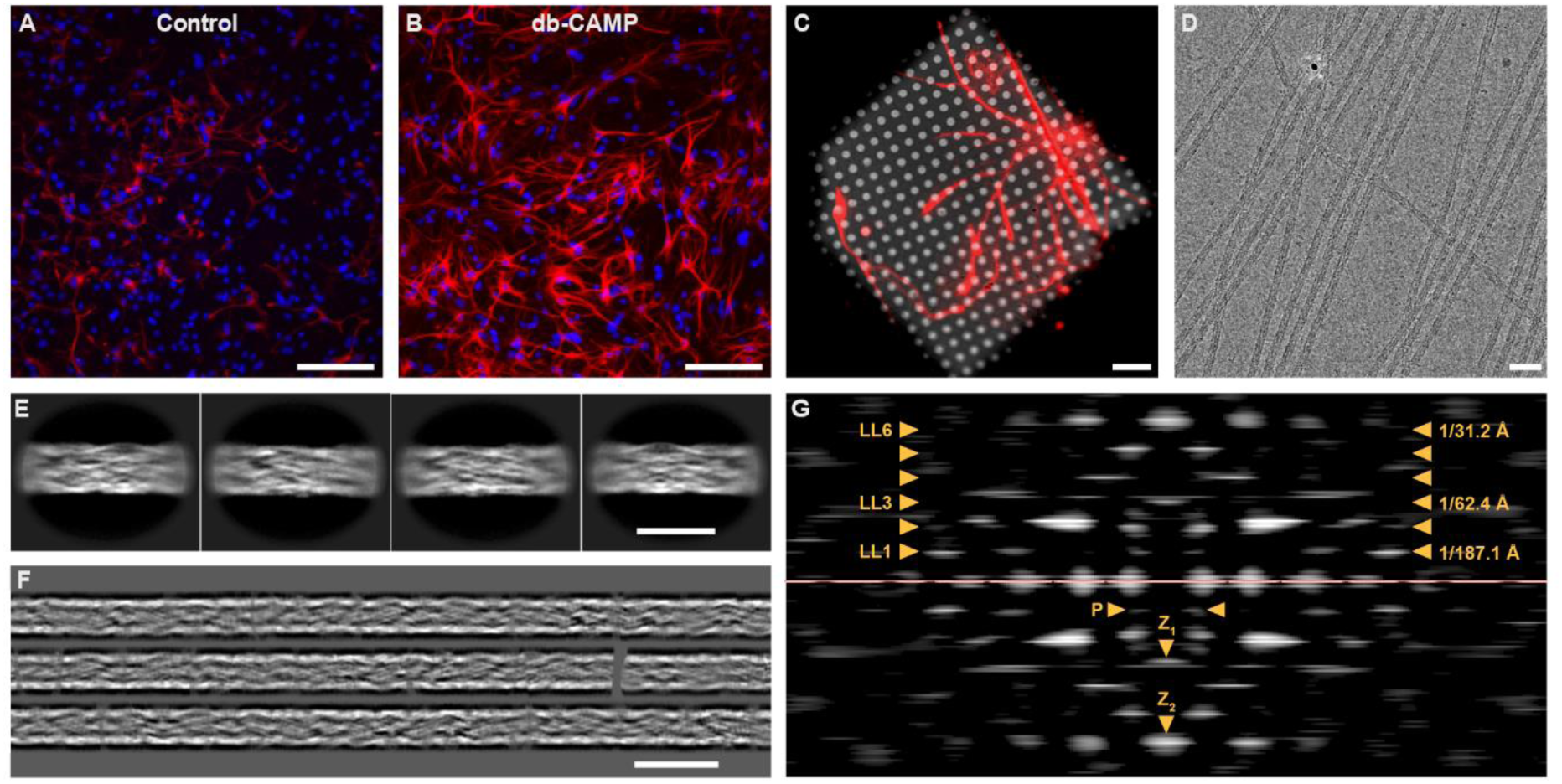
GFAP assembles into helical filaments within astrocytes. **A:** Immunofluorescence of single passage primary *Vim^−/−^* astrocytes. GFAP expression is seen throughout the cytoplasm and extended processes of astrocytes. Many cells display minimal GFAP networks. Cells are stained for GFAP (red) and DNA (blue). Scale bar, 200 µm. **B:** Immunofluorescence of passage 1 primary *Vim^−/−^*astrocytes after a single passage and 14-day treatment with 250 µm db-CAMP. GFAP networks are greatly expanded, with cells extending multiple GFAP containing processes. Cells are stained for GFAP (red) and DNA (blue). Scale bar, 200 µm. **C:** Immunofluorescence of 14-day db-CAMP treated astrocyte on EM grid prior to detergent extraction and vitrification. GFAP forms an extended network throughout the cytoplasm. Cells are stained for GFAP (red). Scale bar, 10 µm. **D:** Cryo-EM micrograph of GFAP filaments acquired after detergent extraction. GFAP filaments are visible as ~10 nm diameter filaments which run largely straight through the field of view. Scale bar, 50 nm. **E:** Class averages of four of the most populous classes after 2D classification of GFAP segments (312 Å box size). GFAP exhibits a complex internal structure and lengthwise diameter fluctuations. Scale bar, 156 Å. **F:** Gallery of three ~210 nm long computationally reconstituted GFAP filaments. Class averages were back-mapped and stitched together before filament straightening. Filaments retain a complex internal structure over long distances. Scale bar, 20 nm. **G:** Mean power spectrum of the computationally reconstituted GFAP filaments (n=51). Multiple layer lines (yellow arrowheads, left) confirm the helical nature of GFAP filaments. The first layer line is visible at ~1/187 Å, with meridional peaks observed at ~1/62 Å and ~1/31 Å (yellow arrowheads, right). Relevant peaks from layer lines LL1, LL3 and LL6 are highlighted below the equator (pink line) as P, Z_1_, and Z_2_, respectively.

GFAP filaments were isolated directly from the db-CAMP treated cells cultured on EM grids through a brief detergent extraction procedure immediately prior to vitrification and imaging by cryo-EM. This sample preparation procedure yields intact, in situ polymerised GFAP filaments while depleting membranes and cytoplasmic components^56^. GFAP filaments are visible in the cryo-EM micrographs as elongated fibres of ~10 nm diameter **(Fig. 1D)**. Compared with in vitro polymerised GFAP filaments **(Extended Data Fig. 1A)**, where the filaments appear significantly more curved and tangled, we consistently found that cell-polymerised GFAP filaments remain largely straight and uniform across the field of view, often running parallel to adjacent filaments.

To assess dataset quality and structural preservation of helical polymers after detergent extraction, we analysed residual F-actin present in the micrographs **(Extended Data Fig. 2A)**. We processed these filaments using their previously determined helical parameters^57^, resolving F-actin to a resolution of ~4.3 Å from ~70,000 particles extracted from the detergent-treated astrocyte data **(Extended Data Fig. 2)**.

To determine the unknown helical parameters of GFAP filaments, we extracted cell-polymerised GFAP segments and subjected the particles to exhaustive 2D classification in both CryoSPARC^58^ and Relion^59^. GFAP class averages exhibit a complex appearance, composed of elongated and twisting structures, as well as diameter fluctuations at the filament surface **(Fig. 1E)**. To visualize the long-range order of the filaments and to facilitate helical parameter determination through power spectrum analysis, we computationally reconstituted extended stretches of GFAP filaments **(Fig. 1F)**.

The resulting power spectrum **(Fig. 1G)** contains a clear set of layer lines, providing direct evidence that GFAP is a helical polymer^60^. Bessel analysis and lattice fitting yields an initial estimate of the in situ GFAP helical parameters, with a helical rise of 62.4 Å and a helical twist of 120° **(Extended Data Fig. 3)**. In silico controls confirm that the helical parameters are not distorted by the computational filament reconstitution procedure **(Extended Data Fig. 4)**. Furthermore, similar helical parameters were also obtained from the analysis of in vitro polymerised GFAP filaments **(Extended Data Fig. 1)**.

### The 3D structure of fully-assembled GFAP filaments

After determining an initial helical parameter candidate, we aimed to solve the 3D structure of fully assembled GFAP filaments. Therefore, we subjected cell-polymerised GFAP segments to extensive helical refinement and 3D classifications with local helical parameter searching **(Supplementary Fig. 1)**, yielding refined helical parameters of 63.3 Å helical rise and 127° helical twist. The resulting 3D structure of the GFAP filament with a global resolution of 7.4 Å is primarily composed of intertwining and interdigitating coiled-coil dimers which are wrapped to form the surface of the filament **(Fig. 2A**, **Extended Data Fig. 5A, Supplementary Video 1)**. These densities are readily segmented, enabling the visualisation of an elongated tetramer with a central tetrameric core and dimeric extensions **(Fig. 2B)**. Overall, this GFAP tetramer adopts an anti-parallel, half-staggered arrangement of coiled-coil dimers, ~65 nm in length, with ~30 nm overlap.

**Fig. 2:**
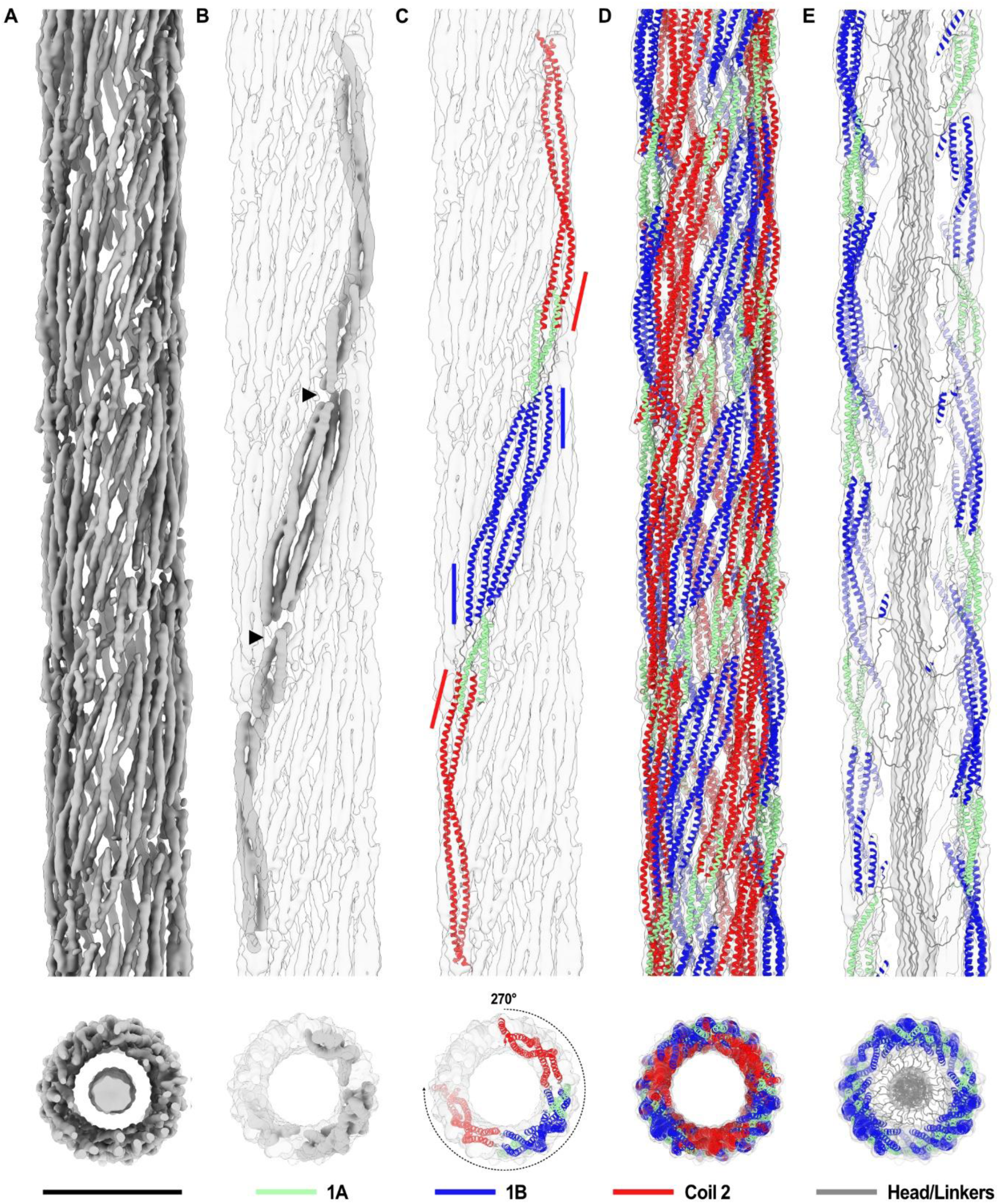
The 3D structure of fully-assembled GFAP filaments. **A:** Density map of the GFAP filament after helical refinement and extension of the map through symmetry imposition. Multiple coiled-coil densities are visible throughout the map. Lower panel: Filament cross-section view of a composite density map containing the luminal fiber low-pass filtered to 30 Å for visualisation. Scale bar, 10 nm. **B:** Segmentation of the GFAP tetramer (dark grey) from the density map (transparent grey). Arrowheads indicate the position of flexible linker domains, which appear as gaps at the chosen isosurface threshold. Lower panel: View along the helical axis illustrating that the tetramer is wrapping around the surface of the filament. Luminal density omitted for clarity. **C:** Model of the GFAP tetramer docked into the density map. The tetramer is composed of anti-parallel dimers, forming a tetrameric core of 1B domains (blue) with overhanging 1A (light green) and Coil-2 domains (red). Blue lines indicate the position of 1B CTEs, red lines the position of Coil-2 NTEs. Lower panel: View along the helical axis of the docked tetramer model, which wraps ~270° around the filament surface. **D:** Model of the GFAP filament tube after symmetry imposition to the tetramer model (63.3 Å rise, 127° twist). Lower panel: View along the helical axis of the filament model. **E:** Cutaway view of the GFAP filament model docked into a composite density map containing the luminal fiber low-pass filtered to 30 Å. Head domains (grey) are modelled to protrude into the filament lumen, forming the highly flexible luminal fiber. Lower panel: View along the helical axis of the filament model with the head domains aggregating into the luminal fiber. Coil-2 omitted for clarity.

Compositional heterogeneity^61^ and local resolution^62^ analyses show that the coiled-coil regions in the density map exhibit high structural homogeneity, as measured by relative occupancy **(Extended Data Fig. 5B)**, and hence exhibit the highest local resolutions in the map **(Extended Data Fig. 5C)**. In contrast, non α-helical regions are marked with lower relative occupancy and are less resolved, suggesting that these densities correspond to flexible, less structurally ordered regions. Amplification^61^ of these flexible densities indicates that well-resolved coiled-coil dimers are connected by short stretches of less-resolved density, likely corresponding to linkers L1 and L12 **(Extended Data Fig. 5B, left; Supplementary Video 2)**. Flexible densities are also observed emanating from the coiled-coil terminals, likely corresponding to parts of the intrinsically disordered head and tail domains. These densities appear to enter the filament lumen and emanate from the filament surface, respectively. The lumen of the filament contains an elongated flexible density **(Fig. 2A**, **Extended Data Fig. 5B, right; Supplementary Video 2)**. This luminal fiber is connected to the inner wall of the filament surface through flexible linking densities.

Next, we incorporated AlphaFold^63^ predictions of the GFAP dimer **(Extended Data Fig. 6)**, previously determined crystal structures of the GFAP 1B domain^51^ and molecular dynamics flexible fitting^64,65^ to assemble a model of the GFAP tetramer **(Fig. 2C)**. This anti-parallel tetramer is centred around a tetrameric core of α-helical 1B domains **(Fig. 2C, blue)**, which is inclined relative to the helical axis. Deviation from coiled-coil geometry is present at the C-terminal end (CTE) region of the 1B α-helices **(Fig. 2C, blue lines)**, caused by the hendecad stutter^47,66,67^. This feature is also found in the 1B crystal structure, suggesting a good alignment between our EM density and previous studies^51^.

The 1B tetrameric core is flanked by splayed 1A α-helices **(Fig. 2C, light green)** forming a V-shaped structure in line with previous IF crystal structures^41,68,69^. The Coil-2 domain **(Fig. 2C, red)** forms an α-helical coiled-coil dimer extending away from the 1B tetrameric core. Alteration of its coiled-coil geometry is observed at the N-terminal end (NTE) region of Coil-2 **(Fig. 2C, red lines)**. Here, the α-helices adopt a parallel geometry for ~40 residues, a feature which is in line with previous studies of the VIM Coil-2^70,71^. In addition, flexible densities connecting the α-helical domains **(Extended Data Fig. 5B, left)** facilitated the incorporation of the linker domains L1 and L12 into the tetramer model. Surprisingly, the full tetramer is twisted around the surface of the filament, encompassing ~270° rotation around the helical axis **(Fig. 2C, Supplementary Video 3)**. Applying the helical parameters to the tetramer model reconstitutes the full filament tube **(Fig. 2D)**. Based on the continuation of flexible densities originating from the 1A NTEs **(Extended Data Fig. 5B, right)**, the head domains were modelled to protrude into the filament lumen, where they aggregate to form the luminal fiber **(Fig. 2E, Supplementary Video 4, Supplementary Video 5)**.

To further explore the helical heterogeneity in GFAP filaments, we performed multiple rounds of 3D classification, yielding several populated classes with stable helical rise values between 63-64 Å but with helical twist values varying between 124-128° **(Extended Data Fig. 7A)**. The distribution of the twist classes analysed along the filaments suggests a spatial relationship exists between twist states, with similar twist states found in close proximity **(Extended Data Fig. 7B)**. Subsequent helical refinement of these classes and flexible fitting of the GFAP filament model indicates variable α-helix positioning between low and high twist states **(Extended Data Fig. 7C/D)**, with the filament appearing to vary between a tightly wound and relaxed state **(Supplementary Video 6)**. Comparison of equivalent Cα positions between the different twist states indicates that variability is highest at 1A residues in proximity to linker L1 and Coil-2 CTE residues **(Extended Data Fig. 7E/F)**, suggesting that these regions act as flexible pivot points around which the positioning of the tetramers varies^69,72,73^.

### GFAP filaments assemble through lateral interactions between tetrameric protofilaments

The basic assembly scheme of a GFAP filament requires the interaction of at least 3 successive tetramers to form a tetrameric protofilament **(Fig. 3A)**. The central tetramer (T_1_) presents 2 NTEs and 2 CTEs of the 1A and Coil-2 domains, respectively. These terminals are engaged by upstream (T_2_) and downstream (T_3_) tetramers, which form the interlock or A_CN_ interactions^52,70,74^ **(Fig. 3A, dashed boxes)**. Here, the splayed 1A α-helices **(Fig. 3B, light green)** of one tetramer interdigitate with the CTEs of the Coil-2 domains **(Fig. 3B, red)** of the successive tetramer, thereby forming an elongated protofilament of 4 α-helices in cross-section **(Supplementary Video 7)**. The addition of two flanking tetramers engages all available interlock positions of the central tetramer.

**Fig. 3:**
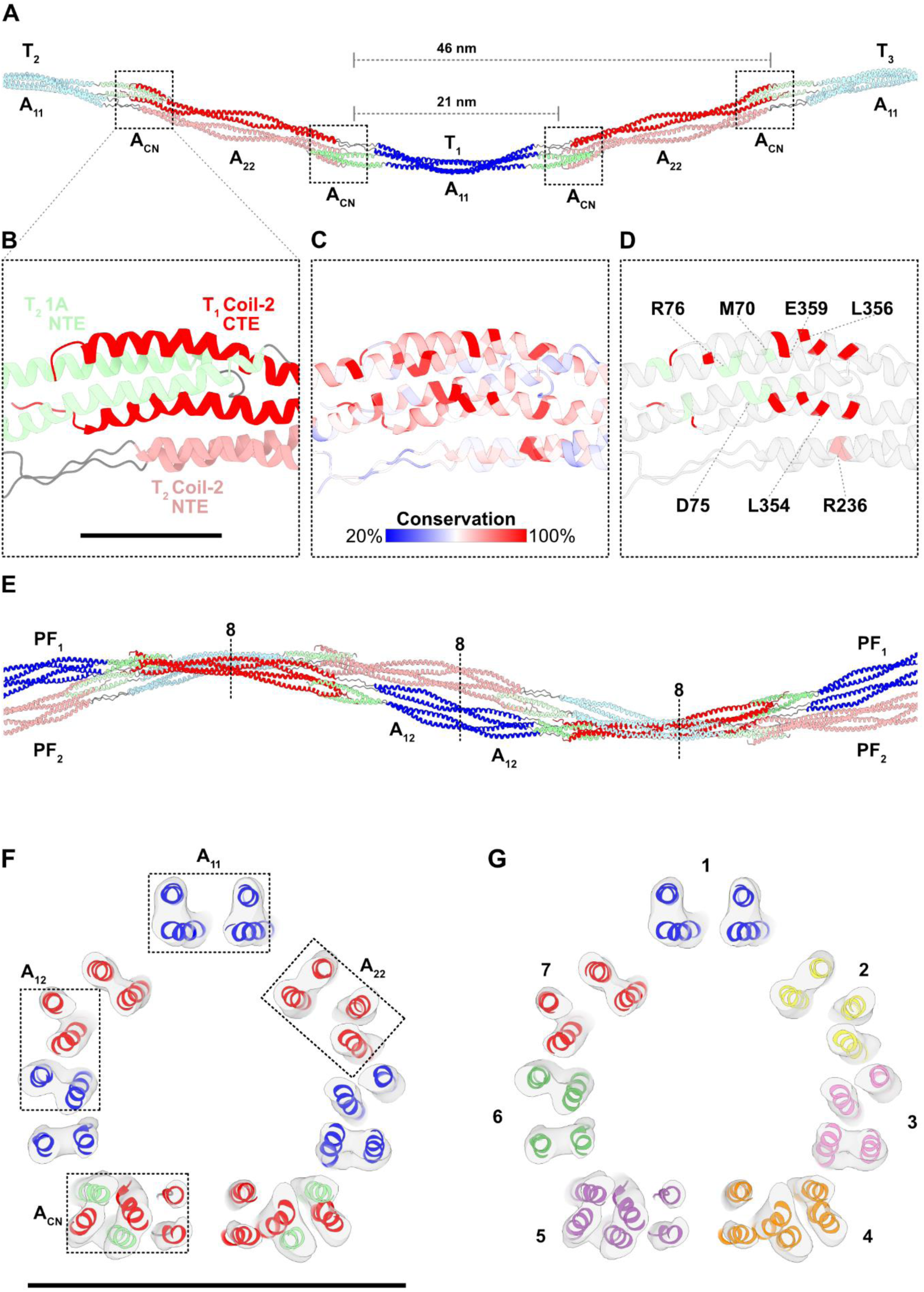
GFAP filaments assemble through lateral interactions between tetrameric protofilaments. **A:** Model of a GFAP protofilament assembled from 3 tetramers. The upstream and downstream tetramers (T_2_/T_3,_ pastel colours) forming 4 interlocks (dashed boxes) with the central tetramer (T_1_, bold colours) through interdigitation of 1A NTEs and Coil-2 CTEs, resulting in the elongation of the protofilament. Labels A_11_, A_22_, A_12_, and A_CN_ refer to the localisation of respective IF binding modes. **B:** Close-up of the interlock formed between the interdigitating 1A NTE of T_2_ (light green) and Coil-2 CTE of T_1_ (red). The linker L12 (grey) and the Coil-2 NTE of T_2_ (light red) run parallel in close proximity to this interaction. Scale bar, 25 Å. **C:** Interlock region coloured according to sequence conservation across multiple IF proteins and families. The interlock region contains the highly conserved residues across the IF family. **D:** Interlock region with positions of mutated residues commonly associated with AxD highlighted (1A NTE: light green, Coil-2 CTE: red, Coil-2 NTE: light red). The labelled residues are most frequently associated with pathology. **E:** Two adjacent protofilaments (PF_1_/PF_2_) forming a protofibril, an assembly with 8 chains in cross-section, which realises the A_12_ binding mode. **F:** The 32-chain cross-section of a fully assembled GFAP filament docked into the density map (transparent grey). All four IF binding modes are present in this cross-section (dashed boxes). **G:** The 32-chain cross-section, color-coded to highlight its 7 constituent protofilaments. The 2 protofilaments with index 4 and index 5 are adding a hexameric cross-section at interlock positions, the other 5 protofilaments adding tetrameric cross-sections.

Additionally, the NTEs of the Coil-2 domains **(Fig. 3B, light red)** are in close proximity to the interlock interaction, suggesting that the parallel arrangement of these α-helices may stabilise the interlock or modulate its functional properties. The arrangement of α-helices in the interlock region creates a short hexameric cross-section spanning ~25 Å, in an otherwise tetrameric protofilament. Intriguingly, the axial distances between the interlocks recapitulate several characteristic features of IFs **(Fig. 3A)**. An axial distance of ~21 nm is seen between proximal interlocks, in line with the periodicity observed by rotary shadowing techniques^43,75,76^, while an axial distance of ~46 nm separates the most distal 1A and Coil-2 interactions in one tetramer, corresponding to previous measurements of IF repeat distances^77,78^.

The interlock region brings the highly conserved residues within the IF family into close spatial proximity **(Fig. 3C)**. Our model indicates that the GFAP interlock hosts the highly conserved residues of both the LNDR motif of domain 1A and the YRKLLEGEE motif of the Coil-2 CTE^68,79,80^, suggesting this interaction is of critical importance for filament assembly and integrity. To understand the importance of this region in human pathology, we mapped mutations associated with AxD pathogenesis to our model^29,40,81^. This reveals that almost all pathogenic mutations are spatially clustered at the filament level, including the mutated R76 and R236 residues^29,82,83^, which are found in the interlock in the 1A LNDR motif and in the adjacent parallel geometry region of the Coil-2 NTE, respectively **(Fig. 3D)**. These two mutations are found in ~37% of all AxD cases^83^.

Visualising the distribution of confirmed pathogenic variants of human GFAP underscores the critical role of interlock and adjacent parallel geometry regions **(Extended Data Fig. 8)**, with almost all pathogenic variants occurring in these areas. These findings provide a molecular basis for the failure of GFAP assembly commonly associated with deleterious AxD variants^27,84,85^ and further highlight the necessity of stable interlock engagement for full GFAP filament assembly. Similar mutations in other IF proteins are also associated with severe pathologies^86–89^, suggesting that the integrity of the interlock region is crucial for IFs in general.

Previous cross-linking experiments have shown that four main binding modes between neighbouring polypeptides are present in fully assembled IFs, referred to as A_11_, A_22_, A_12_, and A ^74,90,91^ interactions. In GFAP filaments, three of these binding modes are formed within a protofilament **(Fig. 3A)**. A_11_ interactions, which involve anti-parallel contacts between 1B α-helices, are intrinsic to the tetramer. A_CN_ interactions arise through the interdigitation of successive tetramers within the interlock. A_22_ interactions occur between anti-parallel Coil-2 domain dimers originating from successive tetramers along the protofilaments.

However, the A_12_ binding mode results from lateral interactions between two protofilaments **(Fig. 3E)**. Such an assembly exhibits 8 chains in cross-section and is termed the protofibril. Based on the GFAP model, this interaction occurs between the 1B CTE hendecad stutter region and the Coil-2 domain of adjacent protofilaments, thereby involving residues ~180-200 and ~280-300, respectively. Interestingly, the latter region contains the only cysteine residue in GFAP, located at position 291. Previous studies have shown that this cysteine is essential for GFAP filament assembly^92^, suggesting that the formation of mature filaments may involve cysteine-mediated interactions between adjacent tetramers.

Our model therefore unifies all four canonical IF binding modes into a helical polymer containing up to 32-chains in cross-section **(Fig. 3F, Supplementary Video 8)**, in line with previously described estimates of IF packing^37^. A fully assembled 32-chain GFAP filament can be described by the placement of 7 protofilaments **(Fig. 3G, Supplementary Video 7)**, with 5 of these protofilaments adding a tetrameric cross-section and 2 adding a hexameric cross-section at interlock positions **(Fig. 3F)**. In total, this assembly requires 11 tetramers to form a ~130 nm-long, minimal-length GFAP filament with a complete 32-chain cross-section **(Extended Data Fig. 9)**.

### GFAP filament bundling in astrocytes

To investigate the higher-order 3D organization of GFAP filaments, we acquired cryo-ET data of db-CAMP treated primary astrocytes, focussing on the cytoplasm of extended processes. Here, we frequently observed tight bundles of straight, parallel and regularly spaced GFAP filaments running through the full field of view of the tomograms **(Fig. 4A, Supplementary Video 9)**. The GFAP bundles were often accompanied by parallel F-actin and came into contact with membrane-bound organelles, including vesicles and endoplasmic reticulum **(Extended Data Fig. 10A)**, suggesting interactions with other cytoskeletal elements.

**Fig. 4:**
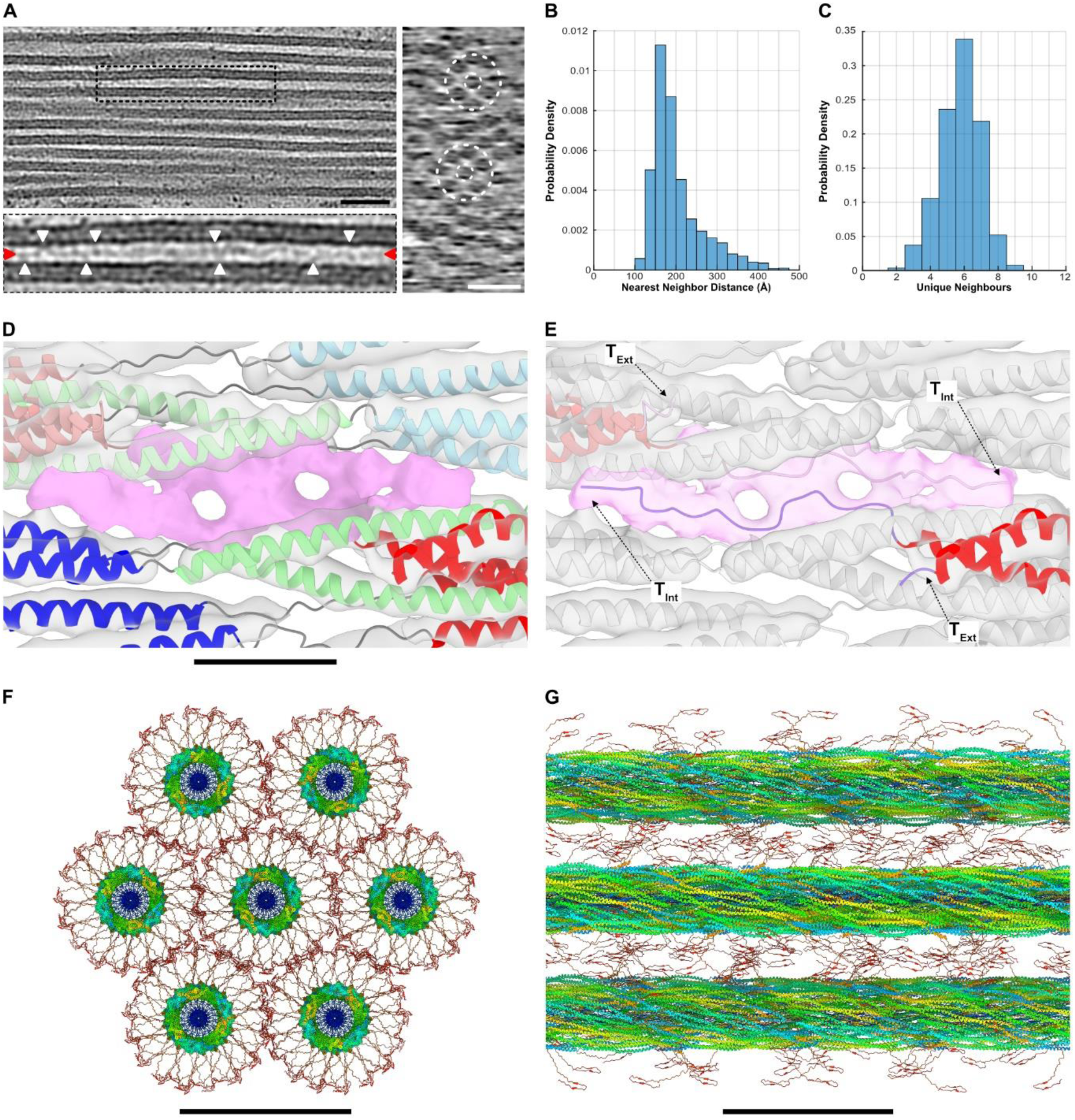
GFAP filament bundling in situ. **A:** Slice (5.4 Å thick) through a GFAP bundle in a tomogram of an db-CAMP treated astrocyte process. The GFAP filaments are oriented parallel to each other in the bundles. Scale bar, 50 nm. Inset: Close-up of the inter-filament space, showing an elongated density (red arrowheads) running between the filaments. Contact points between the inter-filament density and the filament walls are indicated (white arrowheads). Right panel: Slice (yz-plane) through the GFAP bundle, revealing a quasi-hexagonal arrangement of filaments within the bundle core (dashed circles). Scale bar, 50 nm. **B:** Histogram of the nearest-neighbour distances between GFAP filaments (n=419) in GFAP bundles (n=4). The mean nearest-neighbour distance between the filaments is 202 Å ± 62 Å, with a median distance of 183 Å. **C:** Histogram of the neighbour count around each measured filament (n=419). The mean neighbour count is 5.8 ± 1.2, the median is 6. **D:** Close-up of the unassigned density (pink) which is observed at the Coil-2 CTEs (red and pastel red α-helices). The unassigned density is displayed with 50% increased threshold compared to the surrounding map (transparent grey). Scale bar, 25 Å. **E:** Tail model docked into the previously unassigned density. The tails are asymmetrically arranged within each dimer. One tail (T_Ext_) emerges directly from the external surface of the filament and extends outward. The second tail (T_Int_) also projects away from the filament, but only after proceeding for ~16 residues beneath the interlock region, in close proximity to a second, anti-parallel T_Int_ originating from an adjacent tetramer. **F:** Axial view of the GFAP bundle model. Individual filaments are hexagonally arranged with a centre-to-centre spacing of 202 Å. External parts of the tail domains extend ~55 Å from the filament surface to positions equidistant between neighbouring filaments. Tetramers forming the filaments are coloured from blue (N-terminal) to red (C-terminal). Scale bar, 25 nm. **G:** Rotated view of the GFAP bundle model. External parts of the tail domains recapitulate the inter-filament density and contact points observed between the filaments. Scale bar, 50 nm.

Manual tracing of the individual filaments suggests that over 200 filaments can be found in a single GFAP bundle, with bundles with thickness of ~150 nm and width of ~400 nm observed. Nearest-neighbour analysis shows that the mean centre-to-centre distance of the bundled filaments is 202 ± 62 Å **(Fig. 4B)**. Given the ~92 Å diameter of the GFAP model, the surfaces of the filaments are therefore on average separated by ~110 Å within the bundles. Further analysis revealed that bundled GFAP filaments are quasi-hexagonally packed, with filaments often found in proximity to 6 unique neighbours **(Fig. 4A/C**, **Extended Data Fig. 10B)**, in concordance with previous studies of bundled IFs^93–95^.

Close inspection of the inter-filament space in the tomograms revealed an elongated density running between adjacent, in-plane filaments **(Fig. 4A, red arrowheads; Extended Data Fig. 10C)**. This density extends parallel to the filament axes and occupies a position approximately equidistant from the neighbouring filament surfaces. In addition, multiple positions were identified, which show bridging densities between the inter-filament density and the filament surface **(Fig. 4A, white arrowheads)**, suggesting that the inter-filament density could originate from the filaments themselves. Moreover, we observed flexible densities in the cryo-EM reconstruction, which originate from the Coil-2 domain CTEs **(Fig. 4D)** and project outward from the most exterior surface of the filaments **(Extended Data Fig. 5B)**, suggesting that the tail domains are almost completely externalised from the filaments. We therefore speculate that the inter-filament density is a mixture of tail domains originating from neighbouring filaments.

To model the GFAP tail domains, we returned to the cryo-EM density map, identifying unassigned density at the Coil-2 CTE **(Fig. 4D)**. Fitting the tail domains to this density suggests that they are asymmetric within the GFAP dimer **(Fig. 4E)**. One tail (termed T_Ext_) directly emanates from the filament surface at one of the Coil-2 CTEs, while the other tail (termed T_Int_) proceeds beneath the interlock region, before projecting away from the filament. T_Int_ is underpinning the interlock region over a length of ~16 residues. It proceeds in close proximity to the interdigitating 1A NTEs and Coil-2 CTEs and to a second, anti-parallel T_Int_ originating from an adjacent tetramer. Previous work has shown that unlike other IFs, tailless GFAP does not self-assemble into filaments^4,39,96^, suggesting that these rod-proximal tail domain residues may have a role in stabilising or modulating the critical interlock interaction^91^.

Fitting T_Int_ residues into the cryo-EM density map accounts for 32 of the 224 tail domain residues per tetramer, representing ~15% of the total tail length. This suggests that a significant proportion of the tail domains are external and situated around the filaments. In order to include the external tail domains in the GFAP filament model, we added the remainder of the AlphaFold predicted tail domains to the CTEs of T_Ext_ and T_Int_. We placed the conserved β-hairpin motif of the tail^97,98^ **(Extended Data Fig. 6B)** at a distance of ~55 Å from the filament surface, in line with the distance between filament surface and inter-filament density. Finally, we visualised the architecture of a GFAP bundle **(Fig. 4F, Supplementary Video 10)** by arranging the individual filaments hexagonally, with a filament centre-to-centre distance of 202 Å. The accumulation of the tail domains within the inter-filament space recapitulates the inter-filament density observed in the tomograms **(Fig. 4G)**.

## Discussion

Based on cryo-EM and cryo-ET data, we derive a structural model of GFAP filaments in primary astrocytes that reveals an unprecedented molecular architecture of these CNS biopolymers. GFAP filaments are assembled from exceptionally elongated tetramers whose length (~650 Å) exceeds the filament helical rise (63.3 Å) by an order of magnitude. This extreme elongation of the polymer building blocks result in a densely interwoven filament architecture in which multiple tetramers simultaneously traverse individual filament cross-sections. This creates a cross-section of 32-chains in fully assembled GFAP filaments. This assembly integrates all four canonical IF binding modes within a single lattice **(Supplementary Video 8)**. This complex structure can be hierarchically conceptualized based on emerging features observed at the filament level, for example tetrameric protofilaments, octameric protofibrils, and the hexameric interlock region. Such a structural organization stands in sharp contrast to other cytoskeletal polymers^57,99^ and amyloid fibrils^100^, which are typically built from comparatively compact and spatially confined subunits.

Overall, the structure of GFAP filaments is organized into three concentric structural zones. The filament tube is formed primarily by the α-helical coiled-coil domains present within the tetramers. Within the filament lumen, the low-complexity head domains interact to form a central luminal fiber^77,101,102^, whereas the tail domains extend outwards of the filaments into the surrounding cytoplasmic space. These findings establish a structural framework for understanding of how low-complexity regions and regular coiled-coil geometry cooperate to generate the unique properties of individual GFAP filaments and networks^7–10^.

Unveiling the structure of GFAP filaments provides fundamental insight into a protein assembly that is increasingly implicated in neurodegenerative disease^1,103,104^. A central feature of GFAP filaments is the interlock region. Pathogenic mutations associated with AxD spatially cluster in this region. Previous studies have shown that these mutations prevent the assembly of stable, elongated GFAP filaments and result in GFAP aggregations^27,105^. Our structural model suggests that mutations in the interlock region may destabilize GFAP filaments or impair their assembly. In intact GFAP filaments, the head domains are sequestered within the luminal fiber. However, interlock disruption could expose these domains, thereby favouring GFAP aggregation, as observed in patient astrocytes. Our model also suggests that alteration of the cysteine residue could affect the A_12_ binding between adjacent protofilaments, potentially explaining how oxidative stress could lead to GFAP filament disruption. These findings provide a mechanistic link between atomic-scale structural perturbations and astrocytic disease phenotypes.

The GFAP filament structure now enables, for the first time, direct comparison between distinct fully assembled IFs **(Fig. 5)**. The rod domains of murine GFAP and human VIM are highly conserved, with 85% sequence identity at the protein level. Both GFAP and VIM filaments are helical polymers assembled from elongated, anti-parallel tetramers. The tetrameric units of VIM and GFAP are of similar overall length. Both tetramers exhibit an anti-parallel 1B tetrameric core of similar length, with the Coil-2 domain dimers of similar dimensions extending in both axial directions. A major difference is seen in the curvature of the tetramers. The GFAP tetramer is twisted around the filament axis and completes ~270° rotation end-to-end. In contrast, the VIM tetramer is straight and runs parallel to the filament axis^52,106^. These tetramers form the outer surface of both filaments, enclosing a luminal fibre which is linked to the inner surface of the filament tube through unstructured linkers.

**Fig. 5:**
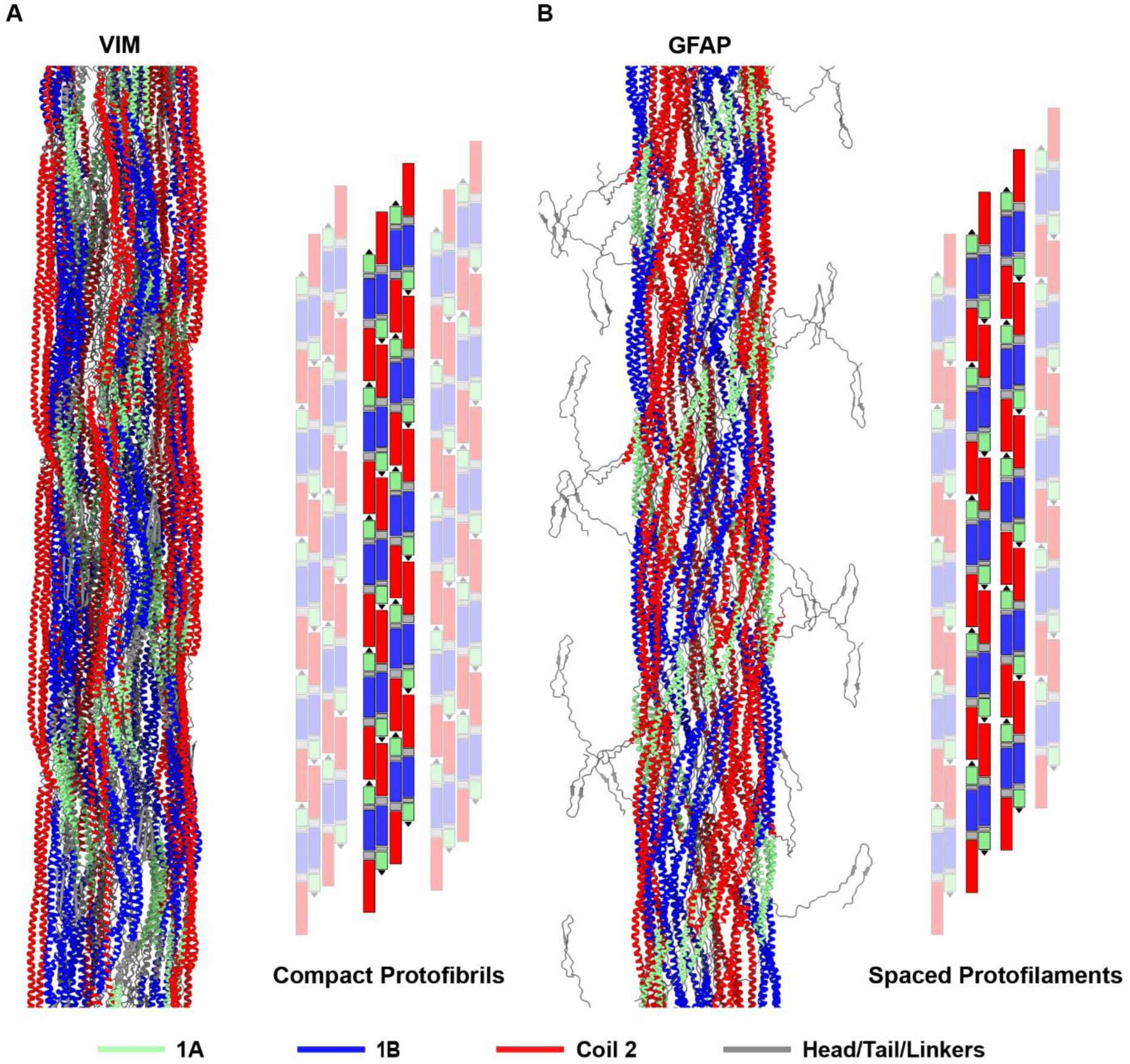
Comparison between VIM and GFAP filaments. **A:** Complete model of VIM filaments. VIM elongates through extension of compact octameric protofibrils running parallel to the helical axis. Protofibrils are linked laterally through the tail domains and centrally through the head domains. Right: lattice representation of VIM assembly. **B:** Complete model of the GFAP filament. GFAP elongates through extension of tetrameric protofilaments, which laterally associate to form octameric protofibrils. Protofibrils laterally associate to form the filament surface and are linked centrally through the head domains. Tail domains are largely extruded from the filament surface. Right: lattice representation of GFAP assembly. Note: Each rectangular component in schematics represents a parallel dimer.

The lumen of VIM filaments is more densely packed than in GFAP filaments, in line with the longer head domain of VIM (85 aa for VIM, 48 aa for GFAP). The tail domains of both proteins are incorporated into the filament periphery to different degrees. In VIM filaments, the tail domain partially fills the inter-protofibril space and facilitates lateral connections between adjacent protofibrils. Although the GFAP tail is largely externalised and possibly involved in inter-filament interactions, a short section of the rod-proximal tail is also found incorporated into the filament, in proximity to the critical interlock interaction. Structural differences between VIM and GFAP filaments indicate that astrocytes may tune IF properties in response to specific stressors through selective upregulation of these proteins.

The incorporation of low-complexity and intrinsically disordered sequences into the filament architecture appears to be a common principle in the assembly of IFs. Visualising flexible and disordered regions remains an outstanding problem in structural biology. In cryo-EM workflows, highly flexible regions such as those in the IFs are often weakly visible or not visible at all as they are averaged out^107^. Consequently, resolution suffers at areas of high flexibility due to misalignment, partial occupancy and subsequent low signal-to-noise ratio^108^. We utilise this property to identify and highlight flexible areas of our map through local resolution and local occupancy estimates. Although we are unable to assign the amino acids and folds in these regions from the electron density map alone, based on location and orientation of the flexible areas in the map we conclude that the terminals of putative 1A and Coil-2 domains take opposite paths. In particular, the 1A α-helices appear to remain associated to the filament tube and inject into the filament core. This suggests that the head domain may contribute to the luminal density, as recently shown for VIM filaments^52^. Analysis of solvent accessibility for the intrinsically disordered segments of IFs suggests that the head domains are indeed buried^109^, while the tail domains show the opposite behaviour. Given the poor local resolution and low occupancy, the luminal density itself is likely highly flexible or exists in a structural continuum. This model is supported by previous experimental evidence demonstrating that isolated low-complexity head domains of IFs, for example the head domain of neurofilament light chain forms amyloid-like fibers^77,101,102^, which are stabilized by transient interactions of cross-β folds.

The externalized tail domains together with the arrangement of filaments in GFAP bundles may result in the formation of a hydrogel-like buffer zone between the bundled filaments counter-acting pressure and ordering the bundles. Assuming an attractive interaction between the β-hairpin motifs in the external tail domains, this would create a multitude of contact points between neighbouring filaments, a hypothesis in-line with the inter-filament density observed in the cryo-ET data. Intriguingly, this architecture would create a buffer zone between the filaments, which maintains inter-filament distance while simultaneously connecting them together. In this way, filament bundle architecture can be stabilized without involving additional cross-linking proteins.

In summary, we present a structural model of GFAP filaments that is consistent with pathological alterations seen in astrocytes. This model provides a framework for designing pharmacological interventions to modulate astrocyte reactivity as a potential treatment strategy in neurological diseases.

## Data availability

The GFAP electron density map is deposited in the Electron Microscopy Data Bank under the accession code EMD-55327. The GFAP model is deposited in the Protein Data Bank under the accession code PDB-9SWW.

## Acknowledgements

This work was funded by grants from the Swiss National Science Foundation (SNSF 320030L-231406) and the Mäxi Foundation to O. M., and from the Swedish Research Council (2024-03159), the Swedish Brain Foundation and EJP RD 2019 ALEXANDER to M. P. We thank the Center for Microscopy and Image Analysis at the University of Zurich.

## Author contributions

C. T. B. conceived the research, prepared the samples, recorded and analyzed all data, developed data analysis methods, built the GFAP model, and wrote the manuscript. M. E. conceived the research, developed data analysis methods, supervised the project and wrote the manuscript. Y. de P. prepared astrocyte cultures and provided feedback on the manuscript. S. B. H. polymerized the in vitro GFAP filaments. N.-H. L. expressed and purified the recombinant GFAP. M.-D. P. guided the GFAP bacterial preparations and provided access to respective resources. M. P. contributed to the project planning and interpretation and edited the manuscript. O. M. conceived the research, supervised the project, edited the manuscript and acquired the funding.

## Materials and Methods

### Animal Handling and Ethics Approval

Mice carrying a null mutation in the vimentin gene (*Vim^−/−^*)^53^ were on a mixed C57Bl6/129 Sv/129Ola genetic background. Mice were housed in standard cages in a barrier animal facility and had free access to food and water. All animal handling and dissection protocols were carried out at the University of Gothenburg. All the experiments were conducted according to protocols approved by the Ethics Committee of the University of Gothenburg.

### Astrocyte Isolation

Astrocyte-enriched cultures were prepared from postnatal day 2 mice as previously described^16^. Mice were decapitated, and brains were dissected under sterile conditions. Cortices were isolated in cold PBS, freed from meninges, and incubated for 10 min at 37 °C in 0.25% trypsin solution. Trypsin was removed, and tissue samples were mechanically dissociated in DMEM supplemented with 10% FBS. The cell suspension was seeded on poly-D-lysine coated T25 flasks. Cells were grown in DMEM supplemented with 10% FBS, 2 mM L-glutamine, 1% penicillin/streptomycin and kept at 37 °C in a humidified atmosphere with 5% CO2. The day after seeding, flasks were vigorously shaken to remove debris and loosely attached non-astrocyte cells, and fresh medium was added. Thereafter, medium was replaced twice a week. Flasks were filled with warm media and sealed for overnight shipment to Zurich.

### Cell Culture

Astrocyte cultures were maintained in DMEM supplemented with 10% FBS, 2 mM L-glutamine, 1% penicillin/streptomycin and kept at 37 °C in a humidified atmosphere with 5% CO2. Cells were washed 3x with pre-warmed PBS before medium was exchanged twice a week. Astrocytes were passaged by washing with pre-warmed PBS and treating with TrypLE (ThermoFisher, 12604013) for 10 minutes, before seeding at 30,000 cells/cm^2^. Astrocytes were maximally passaged once after receipt, immediately prior to experiments.

### db-CAMP Treatment

12 well plates were coated with 1:100 diluted Matrigel (Th.Geyer, 7611631) overnight at 37 °C before washing 3x with sterile PBS. Astrocytes were counted and seeded at 30,000 cells/cm^2^ as described above. Medium was exchanged twice a week until cells reached confluency. db-CAMP (Merck, 28745) was dissolved in water as per manufacturer recommendations and added directly to medium to a final concentration of 250 µM. db-CAMP containing medium was exchanged twice a week. Phase-contrast images were acquired before every medium exchange at 10x magnification using an Olympus IX83 inverted wide-field microscope. Experiment was terminated after 2 weeks treatment by washing cells 3x with pre-warmed sterile PBS before fixing with 3.7% formaldehyde in 10% sucrose at room temperature for 10 minutes. Cells were washed 3x with PBS and stored in PBS at 4 °C until staining.

### Immunostaining

Fixed cells were incubated with 0.1% Triton X-100 in PBS for 10 minutes at room temperature. Cells were washed 3x in PBS for 5 minutes with gentle agitation. Cells were incubated with a blocking solution of 1% BSA, 20 mg/ml glycine in PBS-T (PBS, 0.1% Tween-20) for 1 hour at room temperature. Blocking solution was aspirated before incubation with primary antibodies diluted in 1% BSA PBS-T in a humidified chamber at 4 °C overnight. GFAP (Dako, Z0334) antibody were used at 1:1000 dilution as per manufacturer recommendations. Primary antibody solution was aspirated and cells washed 3x in PBS for 5 minutes with gentle agitation. Cells were incubated with secondary antibodies and 1:10000 Hoechst diluted in 1% BSA PBS-T for 1 hour at room temperature in the dark. GFAP was stained using Cy3 (Lubio, 711165152) and antibody at 1:200 dilution. Cells were washed 3x in PBS for 5 minutes in the dark with gentle agitation and stored in PBS at 4 °C until imaging. Fluorescence images were acquired on an Olympus IX83 inverted wide-field microscope at 10x and 20x magnification before post-processing in Fiji^110^. The above protocol was also applied when imaging cells seeded on EM-grids.

### Expression, purification and assembly of human GFAP-α

For bacterial expression of GFAP, a pET-24d(+) vector containing human GFAP-α was transformed into Rosetta 2(DE3)pLysS (Merck, US171403-3). Bacteria were grown at 37 °C in TBG supplemented with 50 µg/ml kanamycin and 3.4 µg/ml chloramphenicol with shaking at 180 rpm until an optical density of 0.6 was reached. Protein expression was induced by addition of IPTG at a final concentration of 250 µM and grown for 3 hours at 25 °C with 180 rpm shaking. Bacteria were harvested by centrifuging for 15 minutes at 4000 rpm in a pre-cooled centrifuge. Pellets were resuspended in sterile PBS and pelleted at 5000 rpm for 30 minutes. Pellets were flash frozen in liquid nitrogen and stored at −80 °C until further purification.

Overexpressed GFAP presents in inclusion bodies, which were processed as described previously for VIM with minor alterations^111^. In brief, bacterial pellets were sonicated in a buffer containing 50 mM Tris HCl (pH 8), 200 mM NaCl, 25% glycerol, 1 mM EDTA, 10 mg/ml lysozyme, 20 mM MgCl_2_, 8 µg/ml DNase 1, 40 µg/ml RNase A, 1% NP40, 1% deoxycholic acid and one cOmplete Protease Inhibitor Cocktail tablet (Merck, 5056489001) centrifuged at 12,000g for 30 minutes at 4 °C. Inclusions bodies were repeatedly resuspended and centrifuged in a wash buffer containing 10 mM Tris HCl (pH 8), 0.5% Triton X-100, 5 mM EDTA, 1.5 mM DTT and one cOmplete Protease Inhibitor Cocktail tablet until all dark material was removed. A final wash was performed in a buffer containing 10 mM Tris HCl (pH 8), 1 mM EDTA, 1.5 mM DTT and one cOmplete Protease Inhibitor Cocktail tablet. Protein was solubilised with 8 M urea in 10 mM Tris HCl (pH 7.5) and centrifuged at 10,000g for 30 minutes at 4 °C. The supernatant was collected and subjected to size-exclusion-chromatography using a Superdex 200 Increase 10/300 GL column. Protein purity was assessed with SDS-PAGE. Protein fractions were flash frozen and stored at −80 °C until assembly.

Purified GFAP protein was also supplied for testing by Prof. Ming-Der Perng at the National Tsing Hua University in Taiwan. These samples were purified as described in a recent publication^85^. This protocol is largely similar to the above protocol, with an additional anion exchange chromatography step prior to size-exclusion-chromatography.

In vitro assembly was performed as described previously with minor alterations^111^. Briefly, 100 µl of solubilised 0.3 mg/ml GFAP protein was applied to a 12-14 kDa cutoff dialysis device equilibrated with dialysis buffer containing 10 mM Tris (pH 8), 5 mM EDTA, 1 mM DTT. Urea was removed by step wise dialysis in 2 M steps through serial dilution of the solubilisation buffer with dialysis buffer. Buffers were exchanged every 2 hours until an overnight dialysis in urea-free dialysis buffer at 4 °C. Filament assembly was initiated by dialysing into assembly buffer containing 10 mM Tris-HCL (pH 7), 50 mM NaCl, 1 mM DTT for 12-16 hours, heating to 28-30 °C.

Filament assembly was assessed by negative staining transmission electron microscopy. 5 µl of the assembled GFAP solution was applied to holey carbon EM-grids (Cu R2/1, 200 mesh, Quantifoil) and allowed to adsorb for 3 minutes. Excess solution was blotted from the side with filter paper before the grid was rinsed once in distilled water. The grid was stained with 3 µl of 1% (w/v) uranyl acetate for 3 minutes. Excess solution was blotted from the side and the sample air-dried. Micrographs were acquired using a Tecnai G2 Spirit (FEI) operating at 120 kV and processed with Fiji.

### Sample Preparation for Cryo-EM and Cryo-ET

Holey carbon gold EM grids (Au R2/1, 200 mesh, Quantifoil) were glow discharged for 30 seconds using a Harrick-Plasma PDC-32G plasma cleaner at high power before coating with 1:100 diluted Matrigel (Th.Geyer, 7611631) overnight at 37 °C. Grids were washed 3x with pre-warmed PBS before cell seeding. Cell seeding and db-CAMP treatment was conducted as described above. Grids used for cryo-EM of permeabilised samples were rinsed 3x in pre-warmed wash buffer (2 mM MgCl_2_ in PBS) for 30 seconds before immersion in permeabilization buffer (1x PBS, 0.1% Triton X-100, 600 mM KCl, 10 mM MgCl2, 0.02 mg/ml DNase I and protease inhibitors) for 30 seconds. Grids were rinsed 3x in pre-warmed wash buffer and excess liquid removed. Grids were blotted for ~3 seconds from the reverse side before vitrification in liquid ethane using a manual plunge freezing device.

Grids used for cryo-ET of intact cells were washed 3x in pre-warmed PBS for 30 seconds. 3 µl of BSA-coated 10 nm gold fiducial markers (Aurion) was added to the grids immediately prior to blotting and vitrification. Excess liquid was removed from the tweezers before blotting for ~5 seconds from the reverse side before vitrification in liquid ethane using a manual plunge freezing device.

For cryo-EM of in vitro polymerised GFAP, 3 µl of 0.1 mg/ml assembled GFAP was applied to holey carbon EM-grids (Cu R2/1, 200 mesh, Quantifoil) and allowed to settle for 1 minute. Excess liquid was removed before grids were blotted and vitrified using a manual plunge freezing device as described above. Samples were stored under liquid nitrogen for a maximum of 2 weeks until imaging.

### Cryo-ET Acquisition and Tomogram Reconstruction

Grids were loaded into a Titan Krios (FEI) microscope, operating at 300 kV and equipped with a K2 Summit direct electron detector (Gatan) and post-column energy filter (Gatan). First, low magnification maps were acquired at 175x to screen grid quality and identify squares containing cells or cell remnants. Then, medium-magnification maps were acquired at each square containing a cell at 4,800x (calibrated pixel size 6.07 nm/px) to identify suitable positions for tilt-series acquisition. Tilt-series were acquired in zero-loss mode with a slit width of 20 eV using PACEtomo^112^ at a nominal magnification of 81,000x (calibrated pixel size 1.717 Å/px), using a dose-symmetric scheme^113^ from 0° to ±60° in 3° increments with a total dose of ~120 e^−^/Å^2^. Target defocus was set to −4 µm. Acquisition was controlled using SerialEM 4.1^114^.

Frames were processed using the tomotools workflow (https://github.com/tomotools/ tomotools). Briefly, frame series were gain corrected and aligned into tilt images using MotionCor2^115^ before assembly into stacks using the imod package^116^. Stack quality was evaluated manually and poor tilts excluded. Stacks were passed to AreTomo^117^ for initial alignment and image content evaluation. Accepted tilt series were processed using batch mode in imod^118^, with manual adjustments to fiducial tracking. In cases where fiducial tracking was not possible, etomo patch tracking was used. CTF estimation was performed using Ctfplotter^119^ within the imod package. For visualisation, tomograms were binned to 6.868 Å/px before a SIRT-like filter with 15 iterations was applied and the tomogram rotated into XY view. Denoising was applied using cryoCARE^120^ or Topaz^121^. Contrast adjustments, additional rotations and slicing were performed in Fiji or 3dmod.

### Cryo-EM Acquisition

Grids were loaded into a Titan Krios (FEI) microscope, operating at 300 kV and equipped with a K2 Summit direct electron detector (Gatan) and post-column energy filter (Gatan). First, low magnification maps were acquired at 175x to screen grid quality and identify squares containing cell remnants in the case of permeabilised cells or filament aggregations in the case of in vitro prepared samples. Then, medium-magnification maps were acquired at each square containing a target object at 4,800x (calibrated pixel size 6.07 nm/px). These maps were exhaustively manually screened for holes containing abundant, straight and well-spaced filaments. Multiple micrographs were acquired per hole in zero-loss mode with a slit width of 20 eV at a nominal magnification of 105,000x (calibrated pixel size 1.344 Å/px). A defocus range between −0.5 and −3.5 μm was used. Dose-fractionation was used with a frame exposure of 0.1 seconds per frame with the total electron dose per micrograph set to ~60 e^−^/Å^2^. Acquisition was controlled using SerialEM 4.1.

### CryoSPARC Processing

Micrographs were pre-processed with CryoSPARC Live^58^. Movies were pre-processed with patch-based motion correction and CTF estimation using CTFFIND4^122^ and curated based on CTF fit, astigmatism, defocus average, total motion and relative ice thickness. Initial templates were generated by manual picking and subsequent 2D classification of ~5000 particles across the defocus range, before filament tracing and particle extraction with an inter-box distance of 60 Å.

Initial particle curation for both detergent-treated and in vitro polymerised datasets was performed using exhaustive 2D classification in CryoSPARC. Briefly, particles were initially extracted in a box size of 232 pixels (1.344 Å/px). Multiple rounds of 2D classification were applied. Initial uncertainty factor was set to 1 and force max over poses/shifts was disabled until final iterations to separate junk and reduce overfitting. 40 online-EM iterations and 5 final full iterations were used throughout.

To process expanded box filament segments, curated particles were re-extracted in a box size of 522 pixels and binned to 174 pixels (4.032 Å/px) before exhaustive 2D classification. Particles were re-extracted at 522 pixels (1.344 Å/px) for final classifications.

Actin processing was performed using a similar pipeline as described above, with minor alterations. Briefly, actin templates for filament tracing were generated by manual picking before exhaustive 2D classification. Ab initio templates of actin and junk decoys were used to further clean the particle set through heterogeneous refinement. The final particle set was subjected to an asymmetric helical refinement to generate the final template before helical refinement with imposed symmetry (27.6 Å rise, −166.7° twist)^123^. Local CTF correction and particle polishing with Reference Based Motion Correction^124^ were applied to produce the final reconstruction.

### RELION Processing

Curated particles from CryoSPARC were converted for use in RELION 5^125^ by conversion of .cs metadata to STAR file format using the csparc2star command from pyem (https://doi.org/10.5281/zenodo.3576630). STAR files were modified to ensure compatibility with RELION Euler angle conventions and coordinate handling. Briefly, X/Y translations were removed, in-plane (psi) angle was converted to be parallel to the X axis, a 90° tilt prior and randomised rotational angle between −180° and 180° was added. Additional metadata including helical tube ID and helical track length was also added. Particles were re-extracted in RELION in a box size of 232 pixels (1.344 Å/px) before additional cleanup by 2D classifications with T factor of 1 and alignment disabled. A rotationally averaged template was obtained by applying the *relion_reconstruct* command to the STAR file.

For 3D helical refinement and classification, we used the RELION helical toolbox^59^. Particles were initially refined against a rotationally averaged template with power spectrum obtained symmetry imposed (62.4 Å rise, 120° twist, see below). Refinement was repeated against the output average of this job, with local helical searching enabled. Particles were re-extracted in a box of 116 pixels (2.688 Å/px) with translations removed from the STAR files. 3D classification (*k=30*) was performed against a 30 Å low-pass filtered template from initial refinement. Local helical searching was enabled with search bounds of 40-100 Å rise and 110-160° twist around the initial symmetry of 62.4 Å rise and 120° twist. Classes of interest were re-extracted in a box size of 232 pixel (1.344 Å/px) and subjected to iterative helical refinement with local helical searching.

The final reconstruction of GFAP used for model building was generated by helical refinement of the full dataset against a template generated by refinement of the consensus symmetry class (63.3 Å rise, 127° twist). A tight mask was applied, including the outer wall of the filament but excluding the luminal density. Initial angular sampling was set to 0.9°. Psi and tilt angle prior search range was limited to 3°, while random initial rotational angles were imposed. A tau-fudge factor of 4 was applied during refinement. Map was post-processed according to gold-standard criteria and sharpened with an automatically determined global B-factor of −480 Å^2^. OccuPY^61^ was used to amplify and visualise low occupancy and/or low contrast regions of the map. Local resolution was determined using ResMap^62^ and mapped on the OccuPY amplified map in ChimeraX for visualisation^126^.

To generate extended density maps suitable for fitting of a full-length tetramer, reconstructions were re-extracted in a 725-pixel box (1.344 Å/px) and the refined helical symmetry imposed using information from the central 45% of the box. This created a density map 974 Å in length, allowing the fitting of the ~650 Å long tetramer model. Composite maps were generated by displaying the density map of GFAP at a threshold suitable to visualise coiled-coil features alongside a 30 Å low-pass filtered segmentation of the luminal density.

### Visualisation, Segmentation and Model Building

Reconstructed maps were visualised in ChimeraX. Segmentation of reconstructed maps was performed using Segger^127^ within the ChimeraX GUI. Initial dimer models were generated by inputting the FASTA sequence of murine GFAP (P03995, Uniprot) to AlphaFold Multimer^63,128^. The dimer model was trimmed to include only α-helical regions and split into models of 1A/1B or Coil-2, judged as all α-helical regions before and after linker L12, respectively. Initial fitting of the 1B tetramer was performed by rigid body docking of the human 1B tetramer crystal structure^51^ (PDB: 6A9P), followed by replacement of the model by murine AlphaFold 1B dimers in an anti-parallel orientation using MatchMaker^129^. AlphaFold models of the 1A and Coil-2 domains were fit into appropriate densities around the 1B tetrameric core by rigid body docking. L1/L12 linkers, T_Int_ residues and head domains were built into the EM-density manually using ISOLDE^65^. Model was refined by iterating between PHENIX real-space refinement^130,131^, ISOLDE and Coot^132^ with validation reported produced through PHENIX and MolProbity^133^. The final tetramer model was symmetrised using the consensus symmetry and minimised briefly in ISOLDE to resolve symmetry related clashes and form the final filament model. Head domain residues were restrained to the internal surface of the luminal density with the threshold of the EM-density set to 0.002 before release of position restraints and minimising using ISOLDE. External tail domain was modelled by manually bonding the AlphaFold predicted tail domain to the Coil-2 CTE. The predicted β-hairpin motif was maintained by imposing distance restraints between anti-parallel β-strands. The tail domain was positioned manually using ISOLDE at a maximum distance of ~56 Å from the filament surface in order to reflect the inter-filament distance and the potential interaction of tail domains at a point equidistant between aligned and in-plane filaments within bundles. Sidechains were truncated from the model after refinement to reflect the global resolution of the density map.

To visualise the assembly of a protofilament through interdigitation of the 1A NTE and Coil-2 CTE, a helical symmetry of 443.1 Å rise and 169° twist was applied to the tetramer model. This symmetry describes the position of the tetrameric subunit within one protofilament after translations and rotations along the helical axis. Placement of 7 copies of a protofilament through the consensus symmetry recapitulates the full filament. The hexagonal packing of GFAP was visualised by placing filament models on hexagonal vertices spaced by 202 Å in ChimeraX and applying a 60° axial rotation to each model. Comparisons to VIM were performed by applying the published symmetry of the VIM IF^52^ to the deposited VIM tetramer model (PDB-IHM: 9A3R).

All videos were produced by command line scripting in ChimeraX and edited in Adobe Photoshop.

### Computational Filament Reconstitution and Analysis

Filaments were computationally re-assembled using the translations and rotations of class averages as described previously^52,134^ using a custom script in MATLAB R2023a based on functions from the TOM toolbox^135^. In brief, the calculated 2D transformations for each segment (in-plane rotation angle and XY translation) was inverted and applied to associated class averages before mapping to their positions in the micrograph. Class averages were masked to remove background noise outside the filaments. This results in filaments constructed from class averages, with higher signal-to-noise ratios than raw filaments. The filaments were unbent using a MATLAB R2023a script built upon the Fiji straighten function^136,137^. The filament set was filtered to include filaments containing a defined number of segments and a minimum length. A mean power spectrum of the size defined by a minimal filament length of 210 nm was calculated by averaging the power spectra of each individual filament, resulting in a power spectrum with dimensions of 1562×1562 pixels (1.344 Å/px).

To evaluate the effect of inter-box distance on the power spectrum, we re-extracted particles with an inter-box distance of 80 Å in CryoSPARC and processed the segments as described above. The average power spectrum for comparison was formed from reconstituted filaments of equivalent length. To evaluate the presence of long-range order in the reconstituted filaments and evaluate the necessity of computational reconstitution, we permuted the order of class averages mapped to each filament while maintaining the proportions of each class per filament. The average power spectrum of these randomised filaments was constructed as described above. To evaluate contribution of signals to the power spectra originating from the reconstitution approach, we permuted the image content of each class average while maintaining image statistics before reconstitution. The average power spectrum of these filaments containing permuted class averages was constructed as described above.

### Power Spectrum Analysis

Power spectra were analysed with PyHI^138^. Average power spectra from reconstituted filaments were exported from MATLAB. Layer line spacing was set to best intercept the meridional peak repeats. Bessel order fits and helical lattice candidates were evaluated in the GUI by assigning 2 vectors on either side of the meridian. Symmetries were initially screened by comparison of the calculated mass-per-length (MPL) with published data^6^. The MPL was calculated through the equation^139^ MPL = TetramerMass (kDa) / HelicalRise (nm). Candidate symmetries and associated Bessel order assignments were noted and evaluated by iterative helical refinement in RELION 5 as described above. To evaluate the power spectra of single class averages, high quality class averages with box size of 522 pixels (1.344 Å/px) were selected in CryoSPARC and loaded into PyHI. Images were rotated to correct for minor in-plane rotation errors and the contrast adjusted to minimise background in the power spectra. Bessel order fits and helical lattice candidates were evaluated as described above.

### Helical Variability Analysis

3D classification (k=30) with helical searching was performed as described above against a 30 Å low-pass filtered template obtained from helical refinement with consensus symmetry imposed (63.3 Å rise, 127° twist). Local helical searching was enabled with search bounds of 60-68 Å rise and 120-130° twist, with 1Å/1° sampling around an initial symmetry of 63.3 Å rise and 127° twist. The number of particles per class was extracted from the output model STAR file and plotted as a heatmap. Classification indicated variability in helical twist but minimal variability in helical rise. To visualise the spatial relationship between low twist and high twist particles, the particles were back-mapped to their original micrograph and coloured according to their twist state by adaptation of MATLAB functions from the Actin Polarity Toolbox^140^. For visual clarity, particles are classified as low twist or high twist particles, depending on whether their twist angle is below or above the weighted mean twist angle of the full classification, respectively.

Particles from a low twist (123°) and a high twist class (127.3°), each containing >10,000 particles, were re-extracted in a box of 232 pixels (1.344 Å/px). STAR files were edited to remove rotation angle, translation shifts and reset tilt angle to 90°, in order to remove prior helical information stored in the pose of the particles. Particles were then refined against a 15 Å low-pass filtered template obtained from helical refinement with consensus symmetry. No helical symmetry was imposed during refinement. Helical searching was applied post-refinement using relion_helix_toolbox. This search recovered helical parameters of 63.43 Å rise, 123.6° twist and 63.38 Å rise, 127.3° twist for low twist and high twist particles, respectively, which is indicating that these helical parameters are intrinsic to these particles sets.

Refined low and high twist maps were post-processed and symmetry imposed using recovered symmetry values. The GFAP filament model (with head and tail omitted) was docked into each map independently using ISOLDE. The Euclidean distance between each equivalent Cα position in the two models was calculated in ChimeraX using a custom python script. As helical symmetry generates multiple copies of each Cα within the filament, the median Euclidean distance across all symmetry-equivalent Cα positions was used to represent the per-residue displacement between the two models. Cα distance between the low and high twist models was mapped as a blue-white-red colour palette to the original GFAP filament model in ChimeraX. The median Cα distances were plotted as a histogram with a bin width of 5 residues and colour coded according to domain in line with the colour scheme of the tetramer model.

### Filament Packing Analysis

To measure the in situ inter-filament spacing of GFAP filaments from cryo-ET data, the filaments were manually segmented in the tomograms with 3dmod and exported as coordinate sets. Coordinate sets were processed using a custom script in MATLAB R2023a. Briefly, for each GFAP bundle measured (n=4), the filament paths were interpolated by cubic interpolation of points at every pixel (6.868 Å/px), recapitulating the position and orientation of bundled filaments. A convex hull was plotted around the point cloud and shrunk to 75% of the volume towards the centroid, excluding filaments at the edge of the bundle from analysis. Nearest neighbour analysis was performed, by calculating the shortest Euclidean distance for each point to a point on a different filament. Nearest neighbour analysis was restricted to points located within a 48 Å radius along the z-axis. The resulting mean inter-filament distance plus one standard deviation (264 Å) was used to define a 3D search radius to count the number of neighbours surrounding each filament. Data from all measured bundles was combined to generate final plots of inter-filament distance and nearest neighbour counts. The inter-filament distance was used to define the maximum distance of the tail domain from the filament surface, where the diameter of the filament model (~92 Å) was subtracted from the mean inter-filament distance and divided by two, giving the mean point which is equidistant from the surface of two parallel filaments.

### Alexander Disease Mutation Mapping

To visualise the position of AxD associated variants, all variants of human GFAP with confirmed AxD neuropathology or listed as likely pathogenic according to ClinVar classification^141^ were compiled from previously published analyses^40,81^. The equivalent positions on the murine GFAP sequence were identified by sequence alignment using the Clustal Omega algorithm ^142^. The top 3 most prevalent variants found in the interlock regions of 1A NTE and Coil-2 CTE, as well as the common variant residue R236 in the Coil-2 NTE were marked for visualisation. All variants identified in the rod domain were mapped to their respective sub-domains.

## Extended Data Figures

**Extended Data Figure 1:**
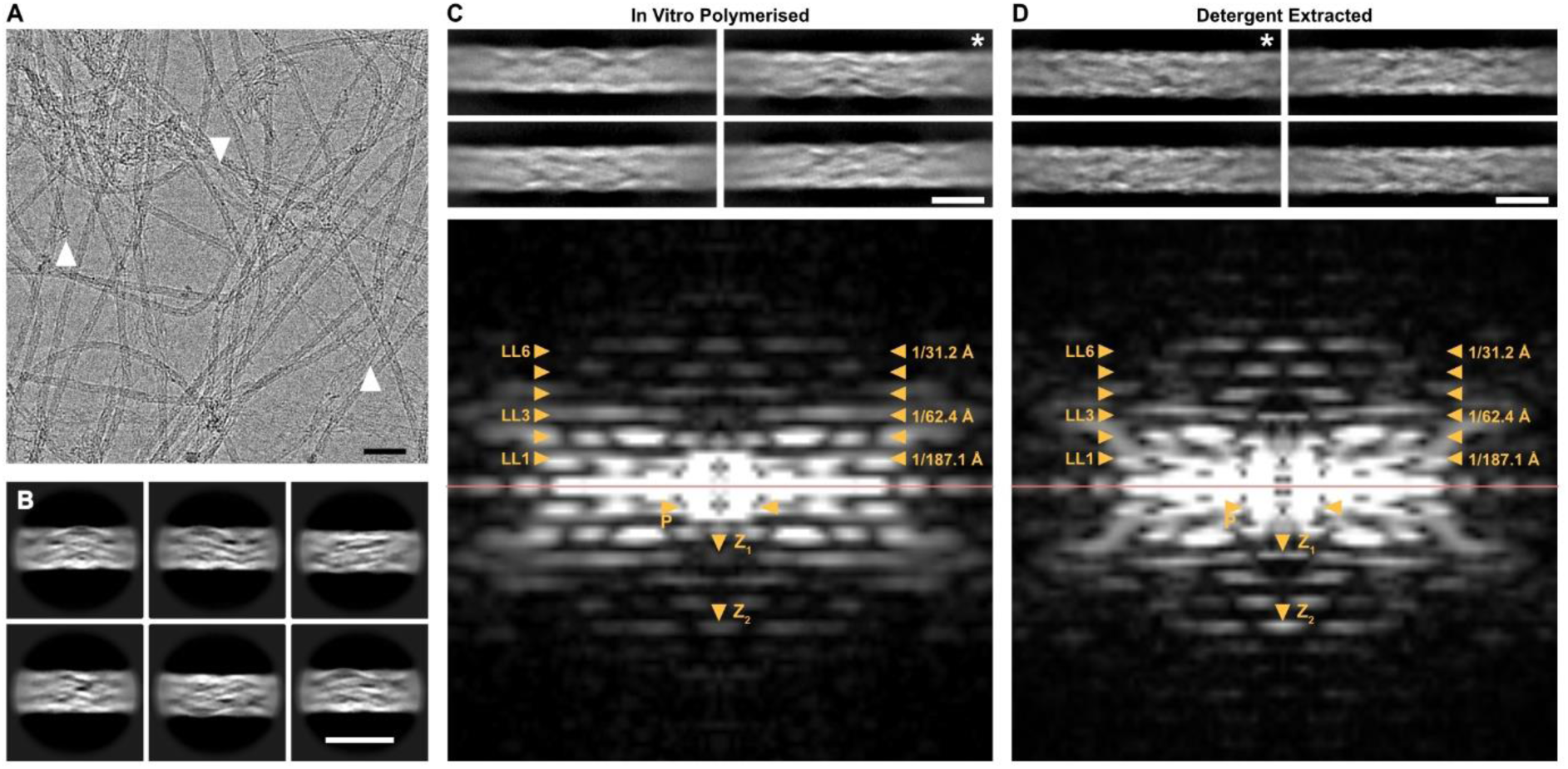
Comparison between in vitro polymerised GFAP filaments and detergent extracted GFAP filaments. **A:** Cryo-EM micrograph of in vitro polymerised GFAP filaments. GFAP filaments are visible as ~10 nm fibres which are often curved and tangled. Unravelling or severing of the filaments can be seen, revealing sub-filament densities (white arrowheads). Scale bar, 50 nm. **B:** Class averages of 6 of the most populous classes after 2D classification of in vitro polymerised GFAP segments (312 Å box size). Similar complex internal structure and diameter fluctuations to cell-polymerised GFAP filaments can be seen. Scale bar, 156 Å. **C:** Class averages of four of the most populous classes after 2D classification of in vitro polymerised GFAP segments in an expanded box (702 Å box size). Some detail can be seen in the centre of the box, with significant blurring towards the box edges. Scale bar, 10 nm. Below: Power spectrum of a single class average (white asterisk). Comparable layer lines and meridional peaks to the montaged power spectrum can be seen (yellow arrowheads). Note the weaker peak at Z_1_ in comparison to the montaged power spectrum and the single class average power spectrum from detergent extracted segments. **D:** Class averages of four of the most populous classes after 2D classification of detergent extracted GFAP segments in an expanded box (702 Å box size). Complex internal structures can be seen throughout the box, with less blurring observed at the box edges. Below: Power spectrum of a single class average (white asterisk). Comparable layer lines and meridional peaks to the montaged power spectrum can be seen (yellow arrowheads). Note the generally increased amplitude for peaks across the spectrum compared to the in vitro polymerised spectrum, especially at Z_1_ and Z_2_.

**Extended Data Figure 2:**
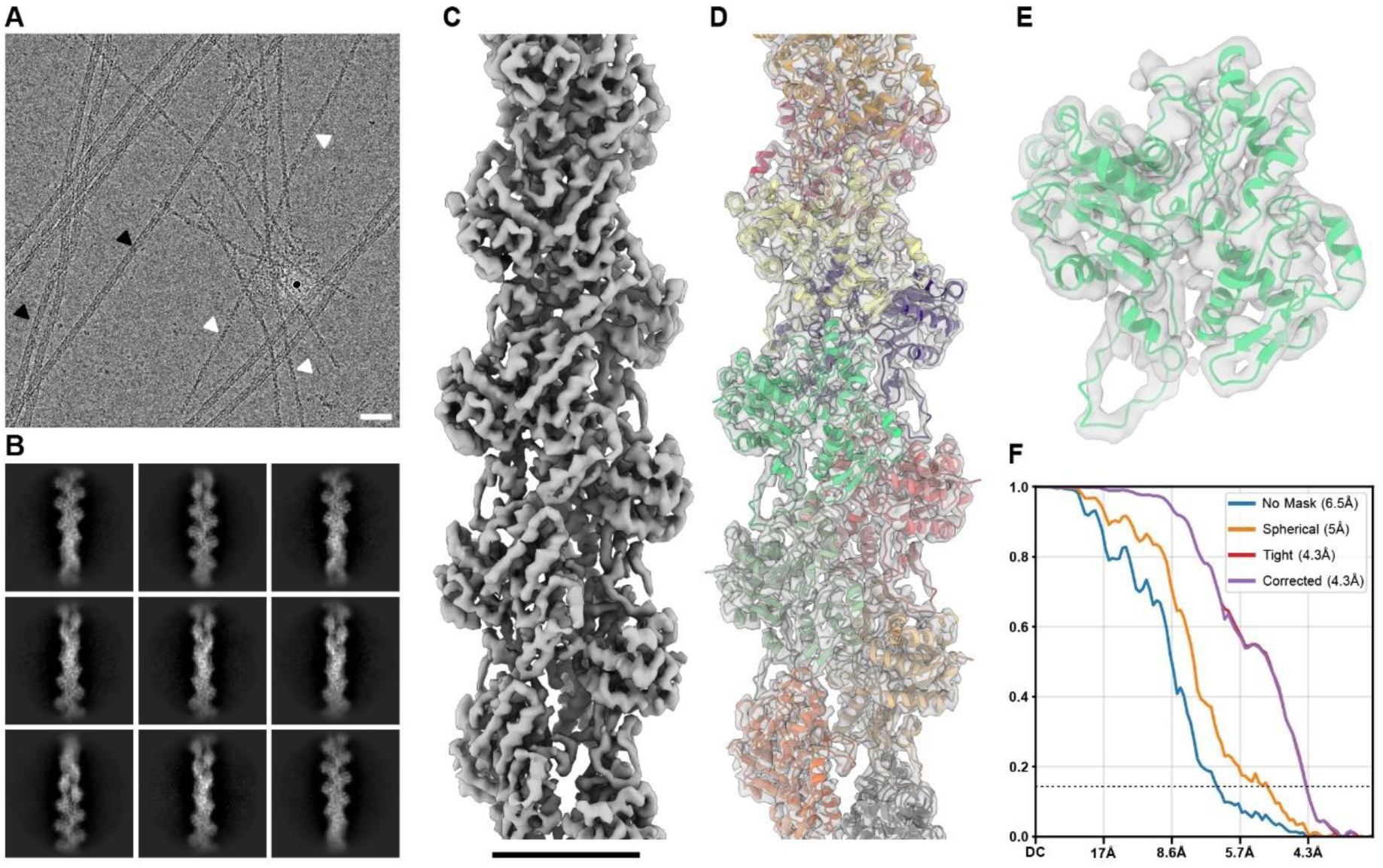
Actin filament refined to ~4 Å after detergent extraction. **A:** Cryo-EM micrograph of GFAP filaments and residual actin filaments acquired after detergent extraction. GFAP filaments are visible as ~10 nm fibres (black arrowheads) and actin filaments as ~7 nm fibres, exhibiting a distinctive zig-zag pattern (white arrowheads). Scale bar, 50 nm. **B:** Class averages of 9 of the most populous classes after 2D classification of actin segments (343 Å box size). **C:** Density map of the actin filament after helical refinement with symmetry imposition (27.6 Å rise, −166.7° twist). Scale bar, 50 Å. **D:** Actin model (PDB: 50NV) docked into the density map (transparent grey) after symmetry imposition. Individual monomers are coloured differently. **E:** Close up of a single actin monomer (PDB: 50NV) fitted into the density of a segmented monomer (transparent grey). **F:** Gold-standard FSC curve of the refined actin structure indicating a global resolution of 4.3 Å at FSC = 0.143.

**Extended Data Figure 3:**
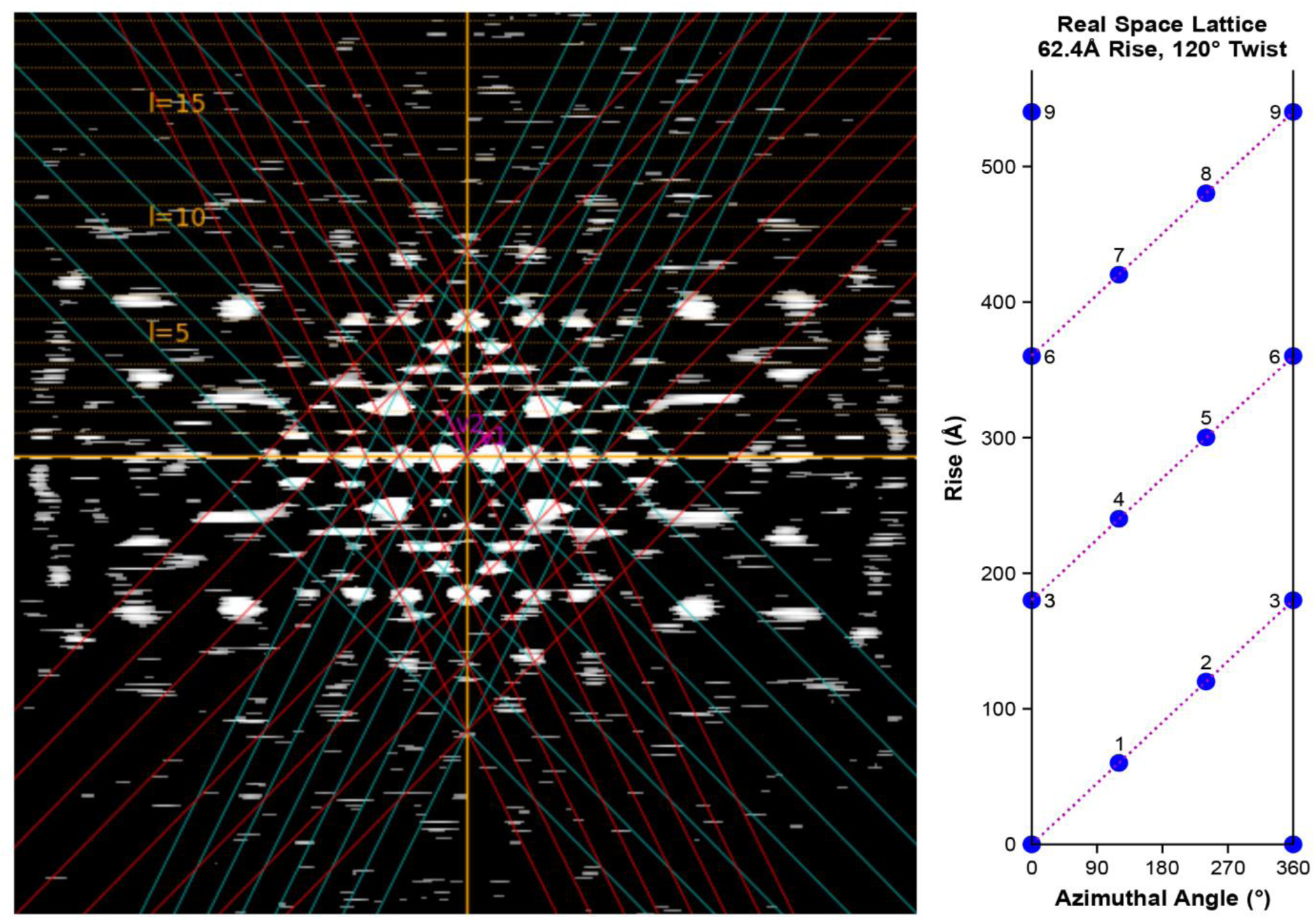
Fourier and real space lattices of power spectrum symmetry candidate. Mean power spectrum of ~210 nm long computationally reconstituted GFAP filaments (n=51) with helical lattice displayed for candidate symmetry 62.4 Å rise and 120° twist. Layer lines are displayed as dashed yellow lines, while fitted vectors 1 and 2 are displayed as blue and red diagonal lines, respectively. Note the intersection of the lattice with most major peaks on the spectrum. Right: Real space lattice describing the position of subunits as defined by the symmetry parameters. There are ~3 subunits placed per 360° rotation.

**Extended Data Figure 4:**
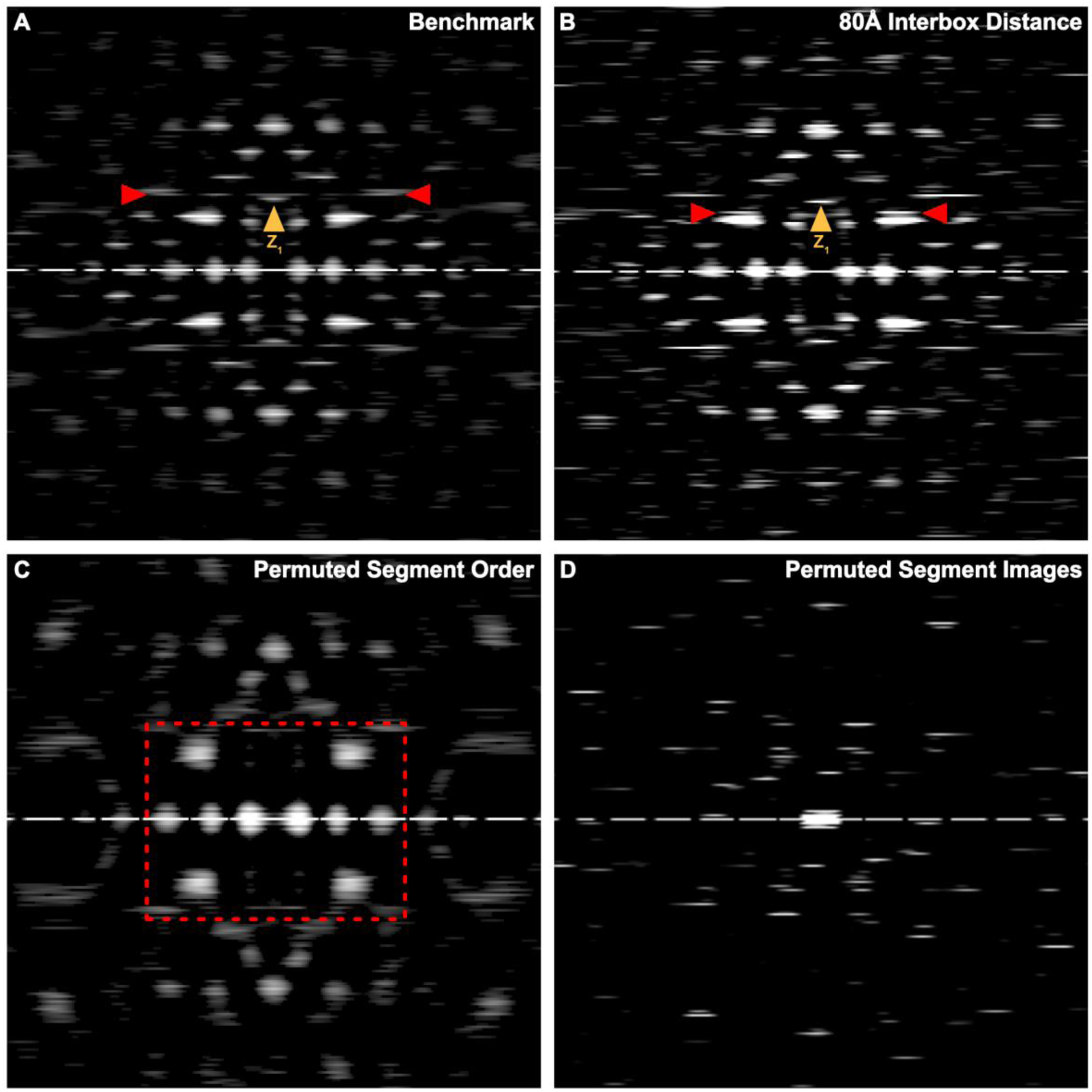
In silico controls of computational filament reconstitution. **A:** Benchmark mean power spectrum of ~210 nm long computationally reconstituted GFAP filaments (n=51) as displayed in Fig. 1G. Red arrowheads mark peaks which originate from the regularly spaced inter-segment distance of 60 Å. The meridional peak at 62.4 Å is marked as Z_1_. **B:** Power spectrum obtained from data with an inter-segment distance of 80 Å. Red arrowheads mark peaks originating from the new inter-segment distance. The 60 Å peaks from the benchmark is absent in this spectrum. The meridional peak at Z_1_ is unaffected. **C:** Power spectrum obtained after permuting the order of class averages along each montaged filament. Many peaks are absent in the central area of this power spectrum (dashed red box). **D:** Power spectrum obtained after permuting the image content of individual class averages.

**Extended Data Figure 5:**
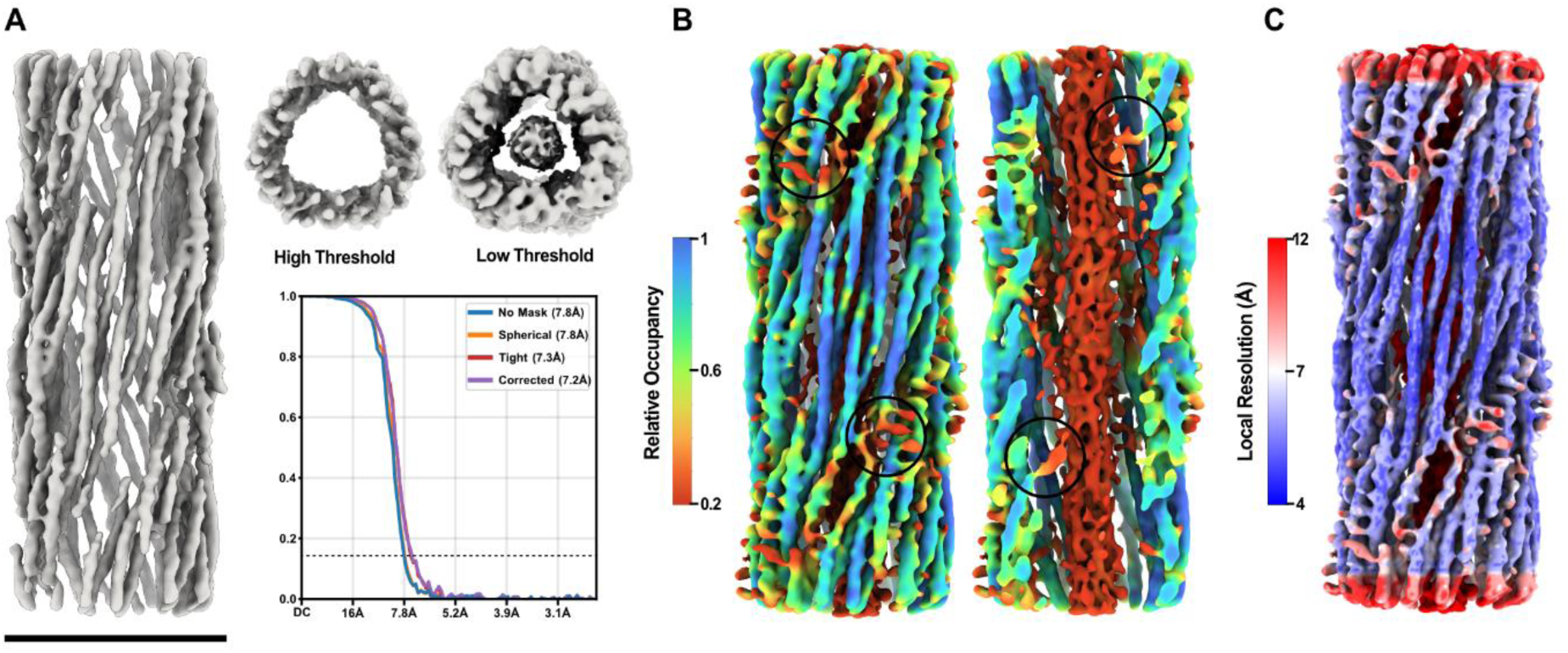
Occupancy and local resolution maps of the GFAP filament structure. **A:** Density map of the GFAP filament (312 Å box size) obtained through helical refinement and symmetry imposition (63.3 Å rise, 127° twist). Upper right: Cross-section views of the GFAP density map at high threshold (~5σ above mean) and low threshold (~2σ above mean), highlighting the appearance of the luminal fiber at low thresholds. Lower right: Gold-standard FSC curve of the refined GFAP structure indicating a global resolution of 7.4 Å at FSC = 0.143. Scale bar, 10 nm. **B:** Amplified density map of the GFAP filament coloured according to relative occupancy as defined by OccuPy^61^. This amplification method allows the visualisation of low signal-to-noise densities while maintaining the overall map sharpness. Flexible regions can be seen as yellow-red densities. Left: Frontal view on the filament tube. Circles indicate positions of flexible linker domains and parts of the tail domains. Right: Cutaway view revealing the filament lumen. Circles indicate connections between the 1A NTEs and the luminal fiber. **C:** Amplified density map of the GFAP filament coloured according to local resolution.

**Extended Data Figure 6:**
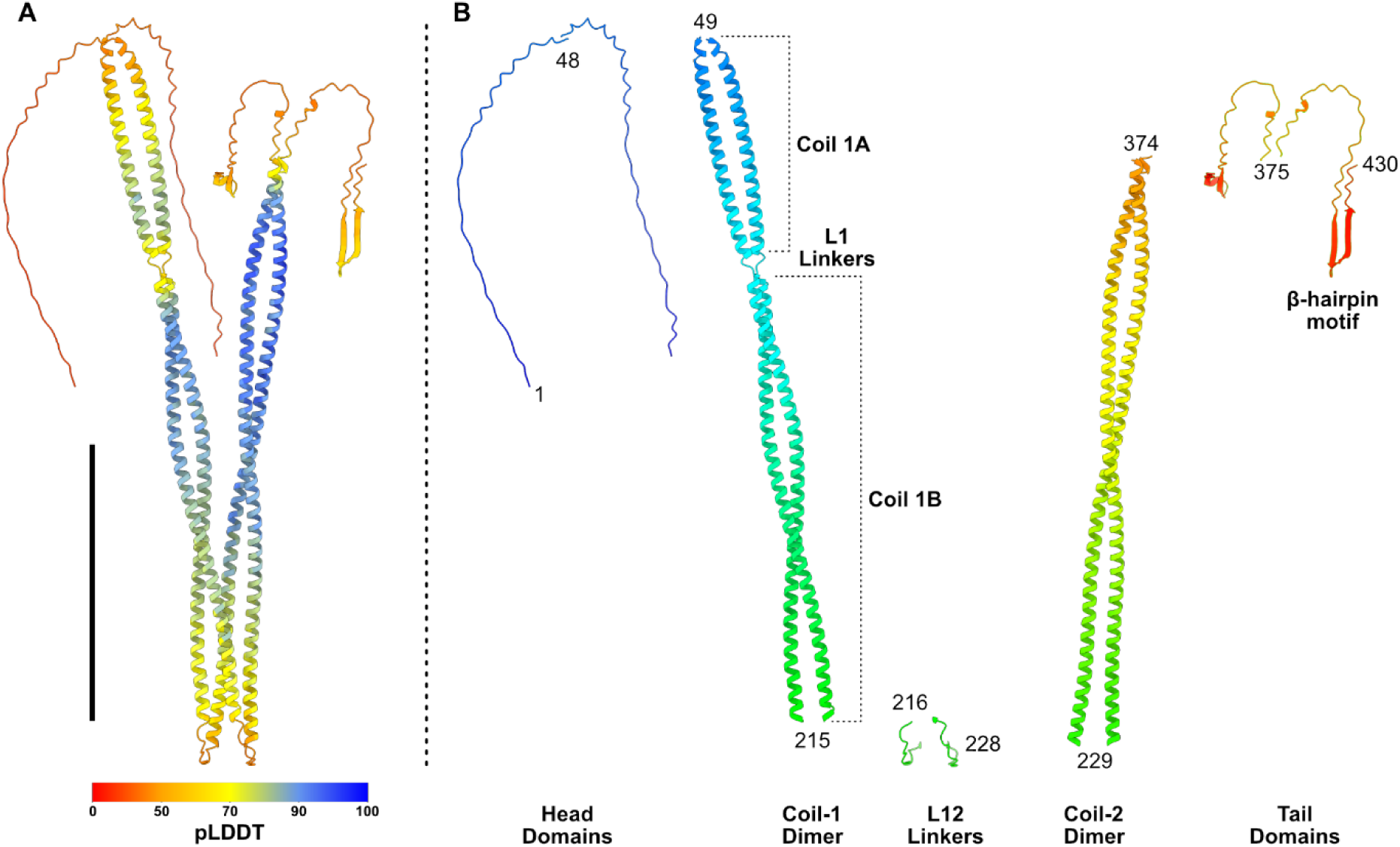
AlphaFold predicted GFAP dimer. **A:** Full murine GFAP dimer model as predicted by AlphaFold. Model is coloured per residue according to the pLDDT value. Scale bar, 100 Å. **B:** Predicted GFAP dimer segmented per domain, coloured from N-terminal (blue) to C-terminal (red). Numbers denote the first and last residue of the labelled domains.

**Extended Data Figure 7:**
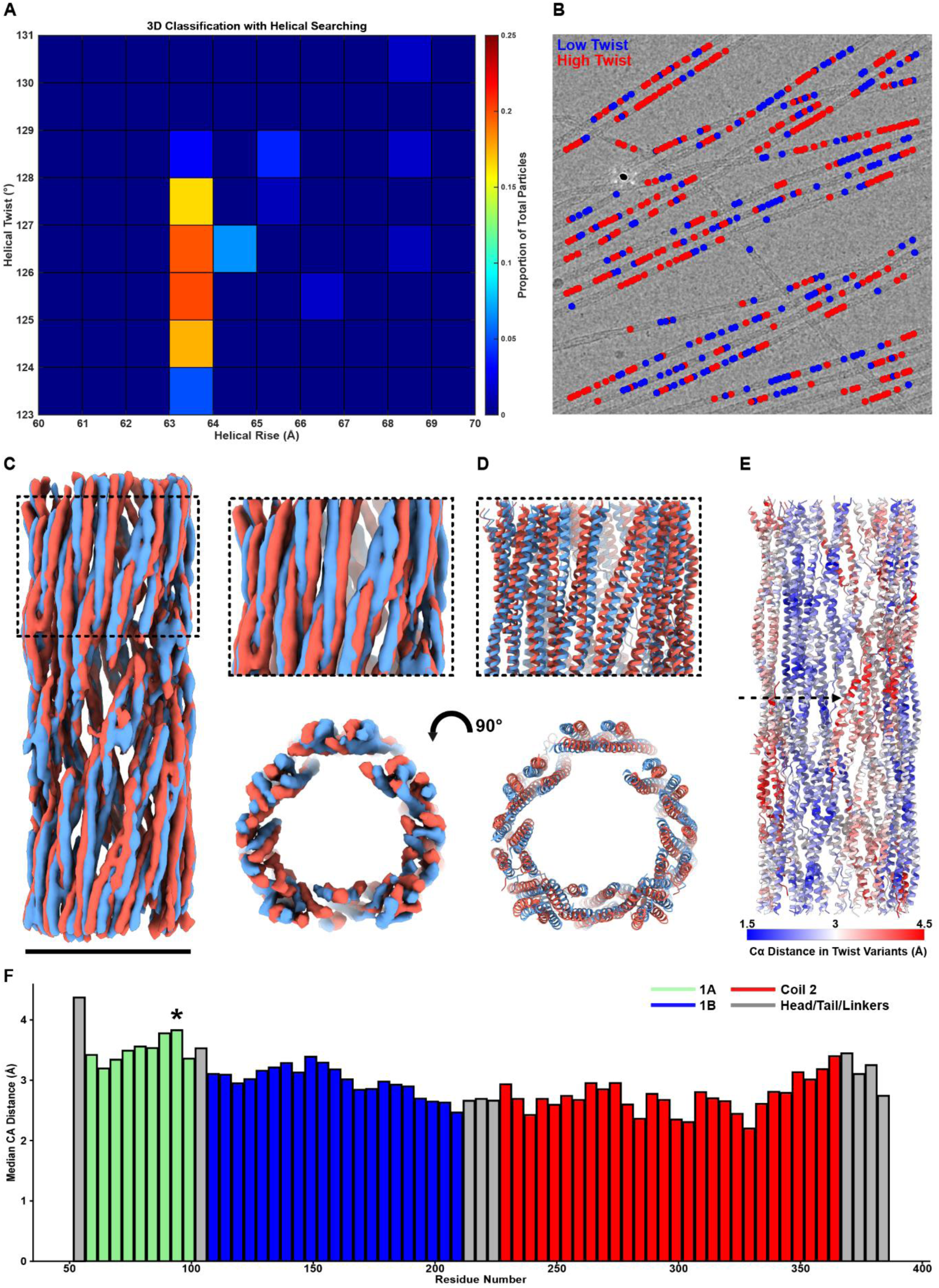
GFAP filaments exhibit continuous helical twist variation along their length. **A:** Heatmap of helical parameter solutions obtained by 3D classification with local helical searches. The majority of particles cluster at a helical rise value between 63-64 Å, but vary in helical twist between ~124-128°. **B:** GFAP segment positions back-mapped to cryo-EM micrograph after 3D classification and assignment to low twist or high twist classes. **C:** Density maps obtained by subsequent helical refinement of low twist and high twist particles obtained through 3D classification. Low twist (blue) and high twist (red) structures were refined with symmetry imposition of 63.43 Å rise, 123.6° twist and 63.38 Å rise, 127.3° twist, respectively. Scale bar, 10 nm. Inset: Close up of the density map highlighting the difference in helix density positioning. Below: Axial view of density maps. **D:** Low twist (blue) and high twist (red) filament models obtained after fitting the consensus structure to the variable twist density maps. Below: Axial view of the models highlighting the difference in helix positions. **E:** Consensus filament model colour coded by the distance between equivalent Cα positions on low twist and high twist models. The maximum movement occurs at residues around the interlock positions (black arrow). **F:** Histogram of median Cα distances for all modelled residues, colour coded according to their domain. The maximum variation occurs at the 1A CTE in proximity to linker L1 (black asterisk) and at the Coil-2 CTE.

**Extended Data Figure 8:**
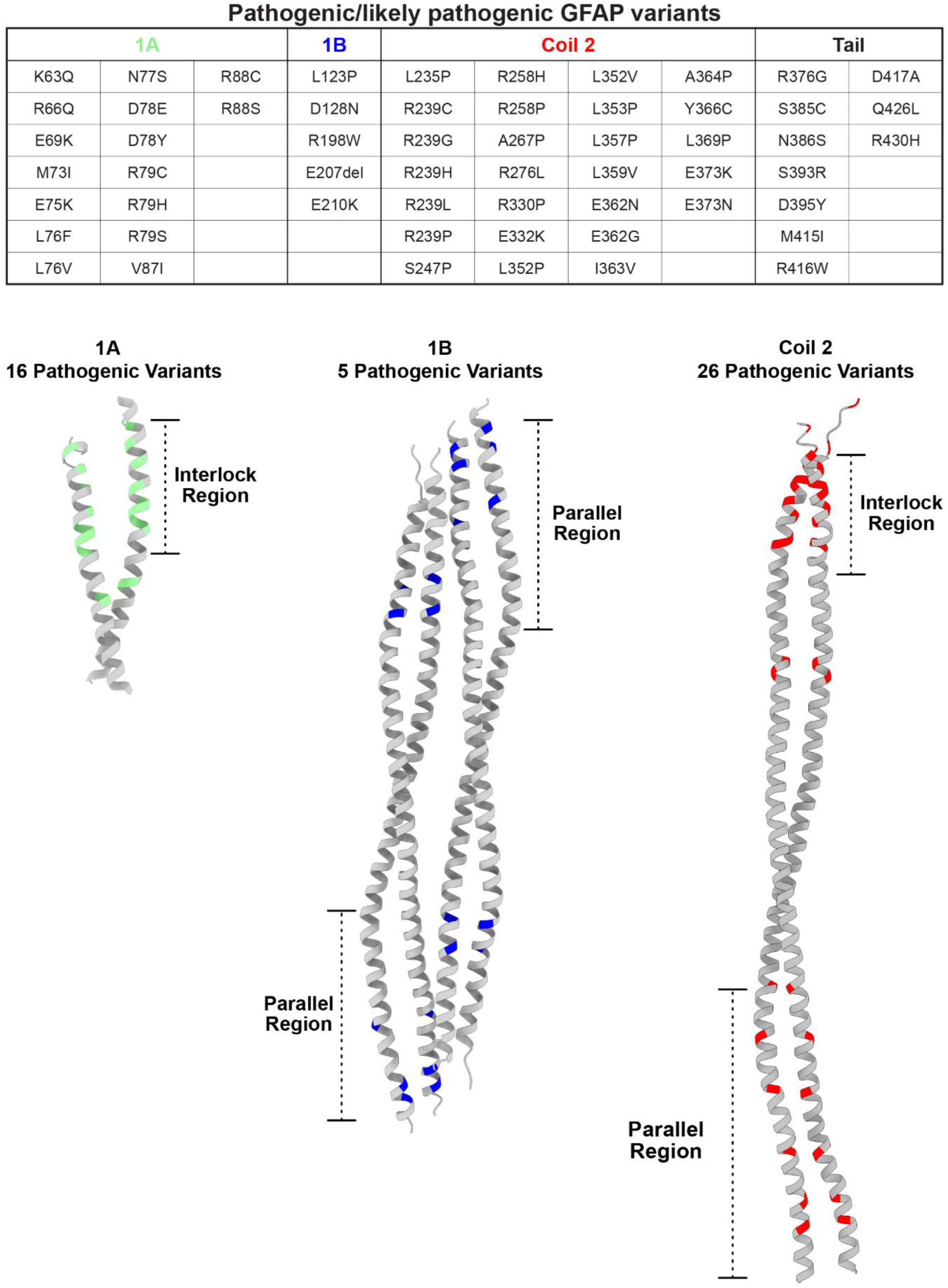
Alexander Disease variants cluster in the interlock and parallel helix regions. **Upper row:** Table of all confirmed pathogenic and likely pathogenic variants of human GFAP associated with AxD. **Below left:** AxD associated variants mapped to the equivalent positions on the 1A domain of the mouse GFAP model. Variants primarily occur in the conserved interlock region at the NTE of 1A. **Below centre:** AxD associated variants mapped to the equivalent positions on the 1B domain of the mouse GFAP model. Variants primarily occur in or in close proximity to the CTE of 1B, in a region where the α-helices run parallel instead of as coiled-coils. **Below right:** AxD associated variants mapped to the equivalent positions on the Coil-2 domain of the mouse GFAP model. Variants are tightly clustered at the interlock region of Coil-2 CTE and distributed across the parallel region of the Coil-2 NTE.

**Extended Data Figure 9:**
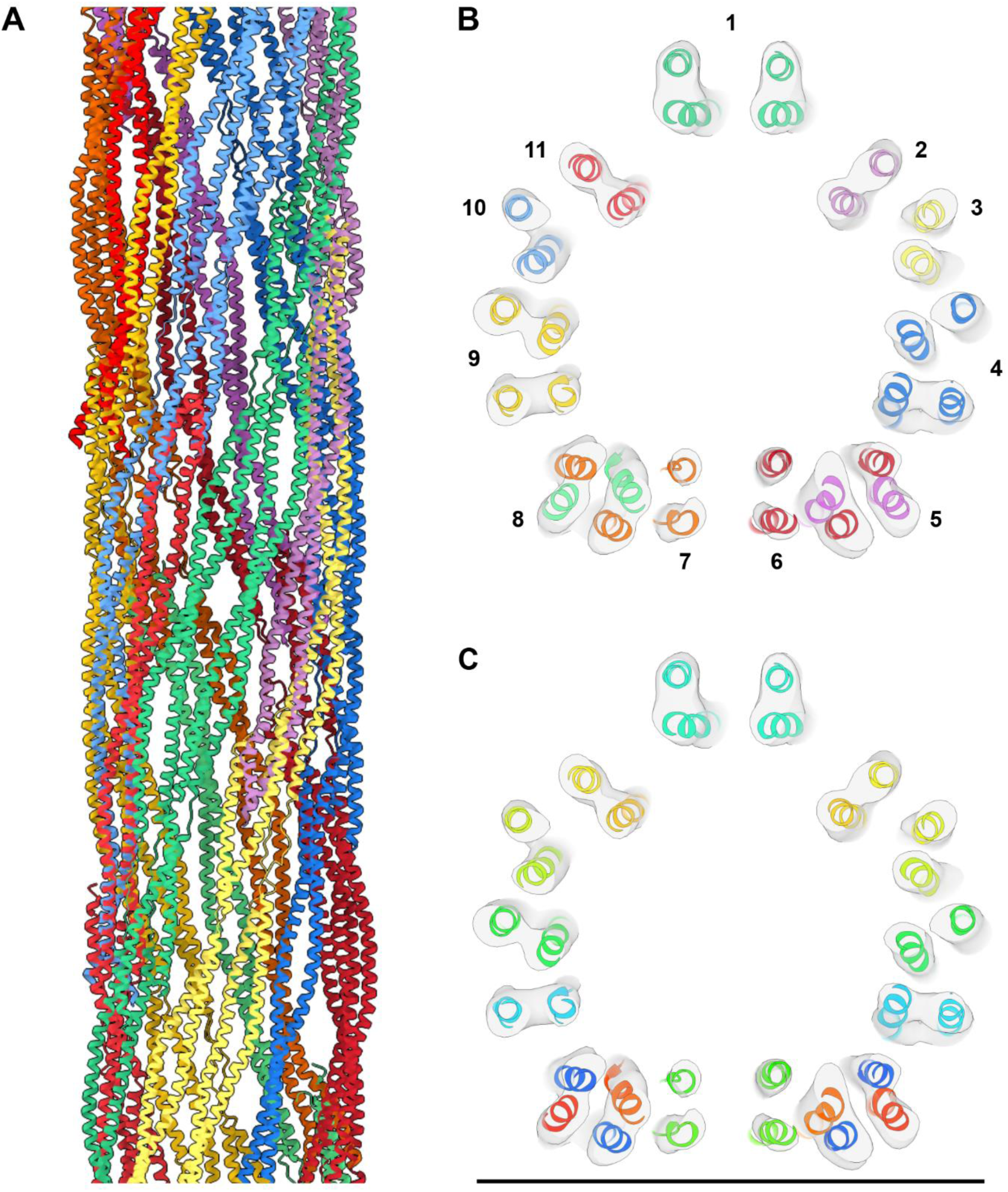
GFAP filaments are composed of 11 tetramers in cross-section. **A:** Side-view of the GFAP filament model with each tetramer coloured individually. **B:** Cross-section view of this model docked into the EM density map (transparent grey). In total, 11 individual tetramers forming the 32-chain cross-section of a GFAP filament. **C:** Axial view of a GFAP filament model with each chain coloured from N-terminal (blue) to C-terminal (red). Scale bar, 10 nm.

**Extended Data Figure 10:**
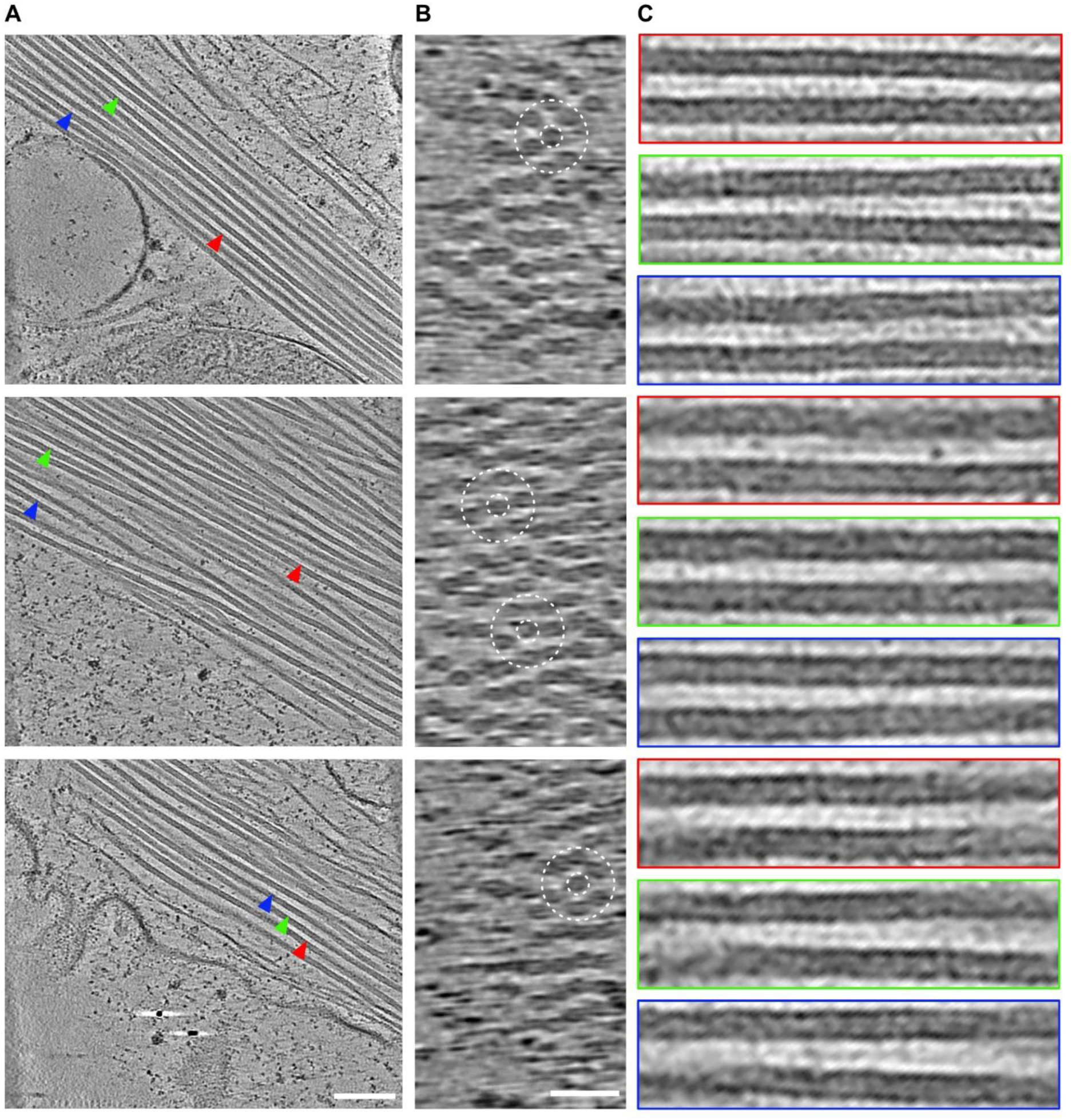
GFAP filaments are highly bundled in situ. **A:** Slices (5.4 Å thick) through GFAP bundles observed in tomograms of extended processes of db-CAMP treated astrocytes. Red, green and blue arrowheads mark positions which are used for close-ups of the inter-filament space. Scale bar, 100 nm. **B:** Slices (yz-plane) through the GFAP bundles. A quasi-hexagonal arrangement of the filaments within the bundle core can be seen (dashed circles). Scale bar, 50 nm. **C:** Rotated close-ups of the inter-filament space extracted from the marked positions in the tomograms. Elongated inter-filament densities are frequently observed, running parallel and equidistant between in-plane filaments.

**Supplementary Figure 1:**
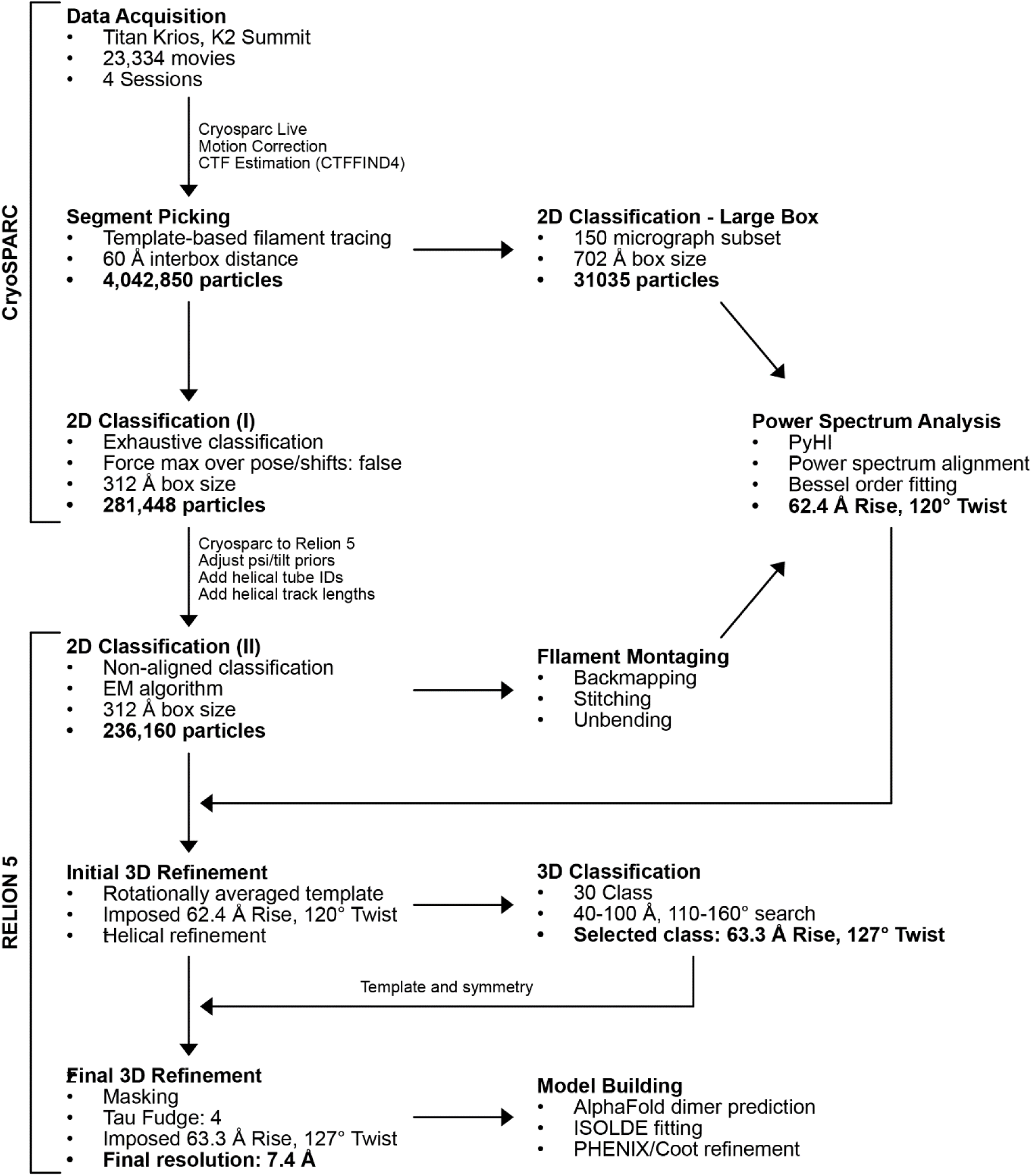
Single particle cryo-EM and helical analysis workflow. Flowchart describing the main processing steps and software used to obtain the 3D reconstruction of the GFAP filament.

## Movie Legends

**Supplementary Video 1:** GFAP density map. This video shows the final refined GFAP structure rotating around the helical axis. A cutaway view with increased threshold to visualise the luminal density is also shown.

**Supplementary Video 2:** GFAP density map amplified with OccuPy and coloured by relative occupancy. This video shows the OccuPy amplified density map of GFAP rotating around the helical axis. A cutaway view is shown to visualise the luminal density and connections to the inner surface of the coiled-coils.

**Supplementary Video 3:** Model of the GFAP tetramer. This video shows the model of the α-helical domains of the GFAP tetramer rotating around the helical axis. The model is shown docked into the extended and symmetry-imposed density map (transparent grey, 63.3 Å rise, 127° twist). The GFAP tetramer wraps ~270° of the filament surface. The 1A domains are shown in light green, 1B domains in blue, Coil-2 in red, and linkers in grey.

**Supplementary Video 4:** Model of the full GFAP filament. This video shows the model of the full GFAP filament rotating around the helical axis. The model is shown docked into the extended and symmetry-imposed density map (transparent grey, 63.3 Å rise, 127° twist). A cutaway view with a composite map including the luminal density low-pass filtered to 30 Å is also shown to visualise the head domain model. The 1A domains are shown in light green, 1B domains in blue, Coil-2 in red, and head domains and linkers in grey.

**Supplementary Video 5:** Axial cross-sections of the full GFAP filament. This video shows cross-sections of the GFAP filament model docked into the density map (transparent grey). The video moves along the helical axis, visualising the relative positions of coiled-coil domains along the filament and displaying the 32 monomer cross section. Threshold is increased through the video to visualise the positioning of the head and linker domains. The 1A domains are shown in light green, 1B domains in blue, Coil-2 in red, and head domains and linkers in grey.

**Supplementary Video 6:** Continuous helical heterogeneity of GFAP filaments. This video shows morphing between the density maps and models of low twist (blue) and high twist (red) structures obtained through 3D classification and helical searching. A morph of refined density maps is shown before the respective low twist and high twist structures are displayed docked into the maps. The filament appears to undergo a breathing motion as it morphs between low twist and high twist states.

**Supplementary Video 7:** Protofilament construction in GFAP filaments. This video shows the successive addition of GFAP tetramers, forming the tetrameric protofilament through fulfilment of the interlock interaction. The addition of 2 tetramers to a central tetramer fulfils all interlocks of the central tetramer. An example of the interdigitation of the 1A NTE and Coil-2 CTE is shown. The combination of 7 tetrameric protofilaments completes the full GFAP filament. The 1A domains are shown in light green, 1B domains in blue, Coil-2 in red, and head domains and linkers in grey. The central tetramer is shown in bold colours, with additional tetramers in light colours.

**Supplementary Video 8:** Emergence of the IF binding modes in GFAP filaments. The video shows the full GFAP filament model, with closeups of the 4 binding modes: A_11_, A_22_, A_CN_, and A_12_. A_11_ is intrinsic to the GFAP tetramer. A_22_ and A_CN_ are emergent from the assembly of protofilaments while A_12_ is emergent from the lateral association of protofilaments in a fully-assembled filament. The 1A domains are shown in light green, 1B domains in blue, Coil-2 in red, and head domains and linkers in grey.

**Supplementary Video 9:** CryoCARE-denoised tomogram of a db-CAMP treated astrocyte process. This video shows successive xy-slices (6.868 Å thick) of a tomogram containing a bundle of GFAP filaments. The tomogram was reconstructed with 4x binning before denoising with a CryoCARE model trained on similar data. Missing wedge correction was not applied. Scale bar, 100 nm.

**Supplementary Video 10:** GFAP filament model with extruded tail domains. This video shows the model of the full GFAP filament with extruded tails rotating around the helical axis. The tail domains extrude to a maximum distance of ~56 Å from the filament surface. The head domain model is visible in the filament lumen. The 1A domains are shown in light green, 1B domains in blue, Coil-2 in red, and head domains, tail domains and linkers in grey.

## References

1 Middeldorp, J. & Hol, E. M. GFAP in health and disease. Prog Neurobiol 93, 421–443 (2011). 10.1016/j.pneurobio.2011.01.005

2 Hol, E. M. & Pekny, M. Glial fibrillary acidic protein (GFAP) and the astrocyte intermediate filament system in diseases of the central nervous system. Curr Opin Cell Biol 32, 121–130 (2015). 10.1016/j.ceb.2015.02.004

3 Kamphuis, W. et al. GFAP Isoforms in Adult Mouse Brain with a Focus on Neurogenic Astrocytes and Reactive Astrogliosis in Mouse Models of Alzheimer Disease. Plos One 7 (2012). https://doi.org/ARTN e42823 10.1371/journal.pone.0042823

4 Lin, N. H., Yang, A. W., Chang, C. H. & Perng, M. D. Elevated GFAP isoform expression promotes protein aggregation and compromises astrocyte function. FASEB J 35, e21614 (2021). 10.1096/fj.202100087R

5 Perng, M. D. et al. Glial Fibrillary Acidic Protein Filaments Can Tolerate the Incorporation of Assembly-compromised GFAP-δ, but with Consequences for Filament Organization and αB-Crystallin Association. Molecular Biology of the Cell 19, 4521–4533 (2008). 10.1091/mbc.E08-03-0284

6 Herrmann, H. & Aebi, U. Intermediate Filaments: Structure and Assembly. Cold Spring Harb Perspect Biol 8 (2016). 10.1101/cshperspect.a018242

7 Lepekhin, E. A. et al. Intermediate filaments regulate astrocyte motility. J Neurochem 79, 617–625 (2001). DOI 10.1046/j.1471-4159.2001.00595.x

8 Yoshida, T. et al. The functional alteration of mutant GFAP depends on the location of the domain: morphological and functional studies using astrocytoma-derived cells. J Hum Genet 52, 362–369 (2007). 10.1007/s10038-007-0124-7

9 Toda, M. et al. Suppression of glial tumor growth by expression of glial fibrillary acidic protein. Neurochem Res 24, 339–343 (1999). Doi 10.1023/A:1022538810581

10 Pekny, M. et al. GFAP-deficient astrocytes are capable of stellation when cocultured with neurons and exhibit a reduced amount of intermediate filaments and an increased cell saturation density. Exp Cell Res 239, 332–343 (1998). DOI 10.1006/excr.1997.3922

11 Wilhelmsson, U. et al. Astrocytes negatively regulate neurogenesis through the Jagged1-mediated Notch pathway. Stem cells 30, 2320–2329 (2012). 10.1002/stem.1196

12 Lebkuechner, I., Wilhelmsson, U., Mollerstrom, E., Pekna, M. & Pekny, M. Heterogeneity of Notch signaling in astrocytes and the effects of GFAP and vimentin deficiency. J Neurochem 135, 234–248 (2015). 10.1111/jnc.13213

13 Wilhelmsson, U. et al. The role of GFAP and vimentin in learning and memory. Biol Chem 400, 1147–1156 (2019). 10.1515/hsz-2019-0199

14 Lundkvist, A. et al. Under stress, the absence of intermediate filaments from Muller cells in the retina has structural and functional consequences. J Cell Sci 117, 3481–3488 (2004). 10.1242/jcs.01221

15 Verardo, M. R. et al. Abnormal reactivity of muller cells after retinal detachment in mice deficient in GFAP and vimentin. Invest Ophthalmol Vis Sci 49, 3659–3665 (2008). https://doi.org/iovs.07-1474 [pii] 10.1167/iovs.07-1474

16 de Pablo, Y., Nilsson, M., Pekna, M. & Pekny, M. Intermediate filaments are important for astrocyte response to oxidative stress induced by oxygen-glucose deprivation and reperfusion. Histochem Cell Biol 140, 81–91 (2013). 10.1007/s00418-013-1110-0

17 Ding, M., Eliasson, C., Betsholtz, C., Hamberger, A. & Pekny, M. Altered taurine release following hypotonic stress in astrocytes from mice deficient for GFAP and vimentin. Brain Res Mol Brain Res 62, 77–81 (1998). 10.1016/s0169-328x(98)00240-x

18 Li, L. et al. Protective role of reactive astrocytes in brain ischemia. J Cereb Blood Flow Metab 28, 468–481 (2008). 10.1038/sj.jcbfm.9600546

19 Wunderlich, K. A. et al. Retinal functional alterations in mice lacking intermediate filament proteins glial fibrillary acidic protein and vimentin. FASEB J 29, 4815–4828 (2015). 10.1096/fj.15-272963

20 Pekny, M. et al. Abnormal reaction to central nervous system injury in mice lacking glial fibrillary acidic protein and vimentin. J Cell Biol 145, 503–514 (1999).

21 Wilhelmsson, U. et al. Absence of glial fibrillary acidic protein and vimentin prevents hypertrophy of astrocytic processes and improves post-traumatic regeneration. J Neurosci 24, 5016–5021 (2004).

22 Cho, K. S. et al. Re-establishing the regenerative potential of central nervous system axons in postnatal mice. J Cell Sci 118, 863–872 (2005).

23 Menet, V., Prieto, M., Privat, A. & Gimenez y Ribotta, M. Axonal plasticity and functional recovery after spinal cord injury in mice deficient in both glial fibrillary acidic protein and vimentin genes. Proc Natl Acad Sci U S A 100, 8999–9004 (2003).

24 Aswendt, M. et al. Reactive astrocytes prevent maladaptive plasticity after ischemic stroke. Prog Neurobiol 209, 102199 (2022). 10.1016/j.pneurobio.2021.102199

25 Pekny, M. & Pekna, M. Reactive gliosis in the pathogenesis of CNS diseases. Biochim Biophys Acta 1862, 483–491 (2016). 10.1016/j.bbadis.2015.11.014

26 Brenner, M. et al. Mutations in GFAP, encoding glial fibrillary acidic protein, are associated with Alexander disease. Nat Genet 27, 117–120 (2001).

27 Yang, A. W., Lin, N. H., Yeh, T. H., Snider, N. & Perng, M. D. Effects of Alexander disease-associated mutations on the assembly and organization of GFAP intermediate filaments. Mol Biol Cell 33, ar69 (2022). 10.1091/mbc.E22-01-0013

28 Der Perng, M., et al. The Alexander disease-causing glial fibrillary acidic protein mutant, R416W, accumulates into Rosenthal fibers by a pathway that involves filament aggregation and the association of alpha B-crystallin and HSP27. Am J Hum Genet 79, 197–213 (2006). 10.1086/504411

29 Quinlan, R. A., Brenner, M., Goldman, J. E. & Messing, A. GFAP and its role in Alexander disease. Exp Cell Res 313, 2077–2087 (2007). 10.1016/j.yexcr.2007.04.004

30 Messing, A. & Brenner, M. GFAP at 50. Asn Neuro 12 (2020). https://doi.org/Artn 1759091420949680 10.1177/1759091420949680

31 Boyd, S. E., Nair, B., Ng, S. W., Keith, J. M. & Orian, J. M. Computational characterization of 3’ splice variants in the GFAP isoform family. Plos One 7, e33565 (2012). 10.1371/journal.pone.0033565

32 Inagaki, M., Nakamura, Y., Takeda, M., Nishimura, T. & Inagaki, N. Glial fibrillary acidic protein: dynamic property and regulation by phosphorylation. Brain Pathol 4, 239–243 (1994). 10.1111/j.1750-3639.1994.tb00839.x

33 Reeves, S. A., Helman, L. J., Allison, A. & Israel, M. A. Molecular cloning and primary structure of human glial fibrillary acidic protein. Proc Natl Acad Sci U S A 86, 5178–5182 (1989). 10.1073/pnas.86.13.5178

34 Mucke, N. et al. Molecular and biophysical characterization of assembly-starter units of human vimentin. J Mol Biol 340, 97–114 (2004). 10.1016/j.jmb.2004.04.039

35 Vermeire, P. J. et al. Molecular Interactions Driving Intermediate Filament Assembly. Cells 10 (2021). 10.3390/cells10092457

36 Kirmse, R. et al. A quantitative kinetic model for the in vitro assembly of intermediate filaments from tetrameric vimentin. J Biol Chem 282, 18563–18572 (2007). 10.1074/jbc.M701063200

37 Herrmann, H., Haner, M., Brettel, M., Ku, N. O. & Aebi, U. Characterization of distinct early assembly units of different intermediate filament proteins. J Mol Biol 286, 1403–1420 (1999). 10.1006/jmbi.1999.2528

38 Herrmann, H. & Aebi, U. Intermediate filament assembly: temperature sensitivity and polymorphism. Cell Mol Life Sci 55, 1416–1431 (1999). 10.1007/s000180050382

39 Chen, W. J. & Liem, R. K. The endless story of the glial fibrillary acidic protein. J Cell Sci 107 (Pt 8), 2299–2311 (1994). 10.1242/jcs.107.8.2299

40 Grossi, A. et al. A systematic review and meta-analysis of GFAP gene variants in Alexander disease. Sci Rep 14, 24341 (2024). 10.1038/s41598-024-75383-4

41 Chernyatina, A. A., Nicolet, S., Aebi, U., Herrmann, H. & Strelkov, S. V. Atomic structure of the vimentin central alpha-helical domain and its implications for intermediate filament assembly. Proc Natl Acad Sci U S A 109, 13620–13625 (2012). 10.1073/pnas.1206836109

42 Qin, Z., Kreplak, L. & Buehler, M. J. Hierarchical structure controls nanomechanical properties of vimentin intermediate filaments. PLoS One 4, e7294 (2009). 10.1371/journal.pone.0007294

43 Parry, D. A. & Steinert, P. M. Intermediate filaments: molecular architecture, assembly, dynamics and polymorphism. Q Rev Biophys 32, 99–187 (1999). 10.1017/s0033583500003516

44 Mucke, N. et al. Assembly Kinetics of Vimentin Tetramers to Unit-Length Filaments: A Stopped-Flow Study. Biophys J 114, 2408–2418 (2018). 10.1016/j.bpj.2018.04.032

45 Parry, D. A., Strelkov, S. V., Burkhard, P., Aebi, U. & Herrmann, H. Towards a molecular description of intermediate filament structure and assembly. Exp Cell Res 313, 2204–2216 (2007). 10.1016/j.yexcr.2007.04.009

46 Kechagia, Z., Eibauer, M. & Medalia, O. Structural determinants of intermediate filament mechanics. Curr Opin Cell Biol 89, 102375 (2024). 10.1016/j.ceb.2024.102375

47 Chernyatina, A. A., Guzenko, D. & Strelkov, S. V. Intermediate filament structure: the bottom-up approach. Curr Opin Cell Biol 32, 65–72 (2015). 10.1016/j.ceb.2014.12.007

48 Guzenko, D., Chernyatina, A. A. & Strelkov, S. V. Crystallographic Studies of Intermediate Filament Proteins. Subcell Biochem 82, 151–170 (2017). 10.1007/978-3-319-49674-0_6

49 Eldirany, S. A., Lomakin, I. B., Ho, M. & Bunick, C. G. Recent insight into intermediate filament structure. Curr Opin Cell Biol 68, 132–143 (2021). 10.1016/j.ceb.2020.10.001

50 Eldirany, S. A., Ho, M., Hinbest, A. J., Lomakin, I. B. & Bunick, C. G. Human keratin 1/10-1B tetramer structures reveal a knob-pocket mechanism in intermediate filament assembly. EMBO J 38 (2019). 10.15252/embj.2018100741

51 Kim, B., Kim, S. & Jin, M. S. Crystal structure of the human glial fibrillary acidic protein 1B domain. Biochemical and Biophysical Research Communications 503, 2899–2905 (2018). 10.1016/j.bbrc.2018.08.066

52 Eibauer, M. et al. Vimentin filaments integrate low-complexity domains in a complex helical structure. Nat Struct Mol Biol 31, 939–949 (2024). 10.1038/s41594-024-01261-2

53 Colucci-Guyon, E. et al. Mice lacking vimentin develop and reproduce without an obvious phenotype. Cell 79, 679–694 (1994). 10.1016/0092-8674(94)90553-3

54 Eliasson, C. et al. Intermediate filament protein partnership in astrocytes. J Biol Chem 274, 23996–24006 (1999). 10.1074/jbc.274.34.23996

55 Le Prince, G., Fages, C., Rolland, B., Nunez, J. & Tardy, M. DBcAMP effect on the expression of GFAP and of its encoding mRNA in astroglial primary cultures. Glia 4, 322–326 (1991). 10.1002/glia.440040310

56 Turgay, Y. et al. The molecular architecture of lamins in somatic cells. Nature 543, 261–264 (2017). 10.1038/nature21382

57 Egelman, E. H., Francis, N. & Derosier, D. J. F-Actin Is a Helix with a Random Variable Twist. Nature 298, 131–135 (1982). DOI 10.1038/298131a0

58 Punjani, A., Rubinstein, J. L., Fleet, D. J. & Brubaker, M. A. cryoSPARC: algorithms for rapid unsupervised cryo-EM structure determination. Nat Methods 14, 290–296 (2017). 10.1038/nmeth.4169

59 He, S. & Scheres, S. H. W. Helical reconstruction in RELION. J Struct Biol 198, 163–176 (2017). 10.1016/j.jsb.2017.02.003

60 Diaz, R., Rice, W. J. & Stokes, D. L. Fourier-Bessel reconstruction of helical assemblies. Methods Enzymol 482, 131–165 (2010). 10.1016/S0076-6879(10)82005-1

61 Forsberg, B. O., Shah, P. N. M. & Burt, A. A robust normalized local filter to estimate compositional heterogeneity directly from cryo-EM maps. Nat Commun 14, 5802 (2023). 10.1038/s41467-023-41478-1

62 Kucukelbir, A., Sigworth, F. J. & Tagare, H. D. Quantifying the local resolution of cryo-EM density maps. Nat Methods 11, 63–65 (2014). 10.1038/nmeth.2727

63 Evans, R. et al. Protein complex prediction with AlphaFold-Multimer. bioRxiv (2021). 10.1101/2021.10.04.463034

64 Trabuco, L. G., Villa, E., Mitra, K., Frank, J. & Schulten, K. Flexible fitting of atomic structures into electron microscopy maps using molecular dynamics. Structure 16, 673–683 (2008). 10.1016/j.str.2008.03.005

65 Croll, T. I. ISOLDE: a physically realistic environment for model building into low-resolution electron-density maps. Acta Crystallogr D Struct Biol 74, 519–530 (2018). 10.1107/S2059798318002425

66 Brown, J. H., Cohen, C. & Parry, D. A. Heptad breaks in alpha-helical coiled coils: stutters and stammers. Proteins 26, 134–145 (1996). 10.1002/(SICI)1097-0134(199610)26:2<134::AID-PROT3>3.0.CO;2-G

67 Parry, D. A. Hendecad repeat in segment 2A and linker L2 of intermediate filament chains implies the possibility of a right-handed coiled-coil structure. J Struct Biol 155, 370–374 (2006). 10.1016/j.jsb.2006.03.017

68 Strelkov, S. V. et al. Conserved segments 1A and 2B of the intermediate filament dimer: their atomic structures and role in filament assembly. Embo J 21, 1255–1266 (2002). DOI 10.1093/emboj/21.6.1255

69 Smith, T. A., Strelkov, S. V., Burkhard, P., Aebi, U. & Parry, D. A. Sequence comparisons of intermediate filament chains: evidence of a unique functional/structural role for coiled-coil segment 1A and linker L1. J Struct Biol 137, 128–145 (2002). 10.1006/jsbi.2002.4438

70 Chernyatina, A. A. & Strelkov, S. V. Stabilization of vimentin coil2 fragment via an engineered disulfide. J Struct Biol 177, 46–53 (2012). 10.1016/j.jsb.2011.11.014

71 Nicolet, S., Herrmann, H., Aebi, U. & Strelkov, S. V. Atomic structure of vimentin coil 2. J Struct Biol 170, 369–376 (2010). 10.1016/j.jsb.2010.02.012

72 Meier, M. et al. Vimentin coil 1A-A molecular switch involved in the initiation of filament elongation. J Mol Biol 390, 245–261 (2009). 10.1016/j.jmb.2009.04.067

73 Ahn, J., Jo, I., Jeong, S., Lee, J. & Ha, N. C. Lamin Filament Assembly Derived from the Atomic Structure of the Antiparallel Four-Helix Bundle. Mol Cells 46, 309–318 (2023). 10.14348/molcells.2023.2144

74 Steinert, P. M., Marekov, L. N. & Parry, D. A. D. Diversity of Intermediate Filament Structure - Evidence That the Alignment of Coiled-Coil Molecules in Vimentin Is Different from That in Keratin Intermediate Filaments. Journal of Biological Chemistry 268, 24916–24925 (1993).

75 Milam, L. & Erickson, H. P. Visualization of a 21-Nm Axial Periodicity in Shadowed Keratin Filaments and Neurofilaments. Journal of Cell Biology 94, 592–596 (1982). DOI 10.1083/jcb.94.3.592

76 Henderson, D., Geisler, N. & Weber, K. A periodic ultrastructure in intermediate filaments. J Mol Biol 155, 173–176 (1982). 10.1016/0022-2836(82)90444-2

77 Lin, Y. et al. Toxic PR Poly-Dipeptides Encoded by the C9orf72 Repeat Expansion Target LC Domain Polymers. Cell 167, 789–802 e712 (2016). 10.1016/j.cell.2016.10.003

78 Vicente, F. N. et al. Molecular organization and mechanics of single vimentin filaments revealed by super-resolution imaging. Sci Adv 8 (2022). https://doi.org/ARTN eabm2696 10.1126/sciadv.abm2696

79 Magin, T. M., Hatzfeld, M. & Franke, W. W. Analysis of Cytokeratin Domains by Cloning and Expression of Intact and Deleted Polypeptides in Escherichia-Coli. Embo J 6, 2607–2615 (1987). DOI 10.1002/j.1460-2075.1987.tb02551.x

80 Herrmann, H. et al. The intermediate filament protein consensus motif of helix 2B: its atomic structure and contribution to assembly. J Mol Biol 298, 817–832 (2000). 10.1006/jmbi.2000.3719

81 Messing, A. Alexander disease associated variants in GFAP, noting which ones have neuropathological confirmation of Rosenthal fibers. (2025). 10.5281/zenodo.17253827

82 Messing, A., Daniels, C. M. L. & Hagemann, T. L. Strategies for Treatment in Alexander Disease. Neurotherapeutics 7, 507–515 (2010). DOI 10.1016/j.nurt.2010.05.013

83 Prust, M. et al. GFAP mutations, age at onset, and clinical subtypes in Alexander disease. Neurology 77, 1287–1294 (2011). 10.1212/WNL.0b013e3182309f72

84 Lin, N. H., Jian, W. S., Snider, N. & Perng, M. D. Glial fibrillary acidic protein is pathologically modified in Alexander disease. Journal of Biological Chemistry 300 (2024). 10.1016/j.jbc.2024.107402

85 Lin, N. H., Jian, W. S. & Perng, M. D. Deletions in Glial Fibrillary Acidic Protein Leading to Alterations in Intermediate Filament Assembly and Network Formation. Int J Mol Sci 26 (2025). 10.3390/ijms26051913

86 Omary, M. B., Coulombe, P. A. & McLean, W. H. I. Mechanisms of disease: Intermediate filament proteins and their associated diseases. New Engl J Med 351, 2087–2100 (2004). DOI 10.1056/NEJMra040319

87 Stone, E. J., Kolb, S. J. & Brown, A. A review and analysis of the clinical literature on Charcot-Marie-Tooth disease caused by mutations in neurofilament protein L. Cytoskeleton 78, 97–110 (2021). 10.1002/cm.21676

88 Cummins, R. E. et al. Keratin 14 point mutations at codon 119 of helix 1A resulting in different epidermolysis bullosa simplex phenotypes. Journal of Investigative Dermatology 117, 1103–1107 (2001). DOI 10.1046/j.0022-202x.2001.01508.x

89 Banerjee, S., Wu, Q., Yu, P., Qi, M. & Li, C. analysis of all point mutations on the 2B domain of K5/K14 causing epidermolysis bullosa simplex: a genotype-phenotype correlation. Mol Biosyst 10, 2567–2577 (2014). 10.1039/c4mb00138a

90 Steinert, P. M., Marekov, L. N., Fraser, R. D. & Parry, D. A. Keratin intermediate filament structure. Crosslinking studies yield quantitative information on molecular dimensions and mechanism of assembly. J Mol Biol 230, 436–452 (1993). 10.1006/jmbi.1993.1161

91 Vermeire, P. J. et al. Molecular structure of soluble vimentin tetramers. Sci Rep 13, 8841 (2023). 10.1038/s41598-023-34814-4

92 Viedma-Poyatos, A., de Pablo, Y., Pekny, M. & Pérez-Sala, D. The cysteine residue of glial fibrillary acidic protein is a critical target for lipoxidation and required for efficient network organization. Free Radical Bio Med 120, 380–394 (2018). 10.1016/j.freeradbiomed.2018.04.007

93 Feughelman, M. & James, V. Hexagonal packing of intermediate filaments (microfibrils) in α-keratin fibers. Text Res J 68, 110–114 (1998). Doi 10.1177/004051759806800205

94 Parry, D. A. D. Structures of the ß-Keratin Filaments and Keratin Intermediate Filaments in the Epidermal Appendages of Birds and Reptiles (Sauropsids). Genes-Basel 12 (2021). https://doi.org/ARTN 591 10.3390/genes12040591

95 Nolting, J. F., Mobius, W. & Koster, S. Mechanics of individual keratin bundles in living cells. Biophys J 107, 2693–2699 (2014). 10.1016/j.bpj.2014.10.039

96 Quinlan, R. A., Moir, R. D. & Stewart, M. Expression in Escherichia coli of fragments of glial fibrillary acidic protein: characterization, assembly properties and paracrystal formation. J Cell Sci 93 (Pt 1), 71–83 (1989). 10.1242/jcs.93.1.71

97 Kouklis, P. D., Papamarcaki, T., Merdes, A. & Georgatos, S. D. A potential role for the COOH-terminal domain in the lateral packing of type III intermediate filaments. J Cell Biol 114, 773–786 (1991). 10.1083/jcb.114.4.773

98 Hess, J. F., Budamagunta, M. S., Aziz, A., FitzGerald, P. G. & Voss, J. C. Electron paramagnetic resonance analysis of the vimentin tail domain reveals points of order in a largely disordered region and conformational adaptation upon filament assembly. Protein Sci 22, 47–55 (2013). 10.1002/pro.2182

99 Nogales, E., Whittaker, M., Milligan, R. A. & Downing, K. H. High-resolution model of the microtubule. Cell 96, 79–88 (1999). Doi 10.1016/S0092-8674(00)80961-7

100 Yang, Y. et al. Cryo-EM structures of amyloid-beta 42 filaments from human brains. Science 375, 167–172 (2022). 10.1126/science.abm7285

101 Zhou, X. M. et al. Transiently structured head domains control intermediate filament assembly. P Natl Acad Sci USA 118 (2021). https://doi.org/ARTN e2022121118 10.1073/pnas.2022121118

102 Zhou, X. M., Kato, M. & Mcknight, S. L. How do disordered head domains assist in the assembly of intermediate filaments? Current Opinion in Cell Biology 85 (2023). https://doi.org/ARTN 102262 10.1016/j.ceb.2023.102262

103 Abdelhak, A. et al. Blood GFAP as an emerging biomarker in brain and spinal cord disorders. Nat Rev Neurol 18, 158–172 (2022). 10.1038/s41582-021-00616-3

104 Li, D. et al. Neurochemical regulation of the expression and function of glial fibrillary acidic protein in astrocytes. Glia 68, 878–897 (2020). 10.1002/glia.23734

105 Jones, J. R. et al. Mutations in GFAP Disrupt the Distribution and Function of Organelles in Human Astrocytes. Cell Rep 25, 947–958 e944 (2018). 10.1016/j.celrep.2018.09.083

106 Eibauer, M. & Medalia, O. Insights into the Structure of Intermediate Filaments. Subcell Biochem 113, 143–161 (2026). 10.1007/978-3-032-05273-5_6

107 Suder, D. S. & Gonen, S. Mitigating the Blurring Effect of CryoEM Averaging on a Flexible and Highly Symmetric Protein Complex through Sub-Particle Reconstruction. Int J Mol Sci 25 (2024). 10.3390/ijms25115665

108 Egelman, E. H. A robust algorithm for the reconstruction of helical filaments using single-particle methods. Ultramicroscopy 85, 225–234 (2000). Doi 10.1016/S0304-3991(00)00062-0

109 Kornreich, M., Avinery, R., Malka-Gibor, E., Laser-Azogui, A. & Beck, R. Order and disorder in intermediate filament proteins. FEBS Lett 589, 2464–2476 (2015). 10.1016/j.febslet.2015.07.024

110 Schindelin, J., et al. Fiji: an open-source platform for biological-image analysis. Nat Methods 9, 676–682 (2012). 10.1038/nmeth.2019

111 Herrmann, H., Kreplak, L. & Aebi, U. Isolation, characterization, and in vitro assembly of intermediate filaments. Methods Cell Biol 78, 3–24 (2004). 10.1016/s0091-679x(04)78001-2

112 Eisenstein, F. et al. Parallel cryo electron tomography on in situ lamellae. Nat Methods 20, 131–138 (2023). 10.1038/s41592-022-01690-1

113 Hagen, W. J. H., Wan, W. & Briggs, J. A. G. Implementation of a cryo-electron tomography tilt-scheme optimized for high resolution subtomogram averaging. J Struct Biol 197, 191–198 (2017). 10.1016/j.jsb.2016.06.007

114 Mastronarde, D. N. SerialEM: A Program for Automated Tilt Series Acquisition on Tecnai Microscopes Using Prediction of Specimen Position. Microscopy and Microanalysis 9, 1182–1183 (2003). 10.1017/s1431927603445911

115 Zheng, S. Q. et al. MotionCor2: anisotropic correction of beam-induced motion for improved cryo-electron microscopy. Nat Methods 14, 331–332 (2017). 10.1038/nmeth.4193

116 Kremer, J. R., Mastronarde, D. N. & McIntosh, J. R. Computer visualization of three-dimensional image data using IMOD. J Struct Biol 116, 71–76 (1996). 10.1006/jsbi.1996.0013

117 Zheng, S. et al. AreTomo: An integrated software package for automated marker-free, motion-corrected cryo-electron tomographic alignment and reconstruction. J Struct Biol X 6, 100068 (2022). 10.1016/j.yjsbx.2022.100068

118 Mastronarde, D. N. & Held, S. R. Automated tilt series alignment and tomographic reconstruction in IMOD. J Struct Biol 197, 102–113 (2017). 10.1016/j.jsb.2016.07.011

119 Mastronarde, D. N. Accurate, automatic determination of astigmatism and phase with Ctfplotter in IMOD. J Struct Biol 216, 108057 (2024). 10.1016/j.jsb.2023.108057

120 Buchholz, T. O. et al. Content-aware image restoration for electron microscopy. Methods Cell Biol 152, 277–289 (2019). 10.1016/bs.mcb.2019.05.001

121 Bepler, T., Kelley, K., Noble, A. J. & Berger, B. Topaz-Denoise: general deep denoising models for cryoEM and cryoET. Nat Commun 11, 5208 (2020). 10.1038/s41467-020-18952-1

122 Rohou, A. & Grigorieff, N. CTFFIND4: Fast and accurate defocus estimation from electron micrographs. J Struct Biol 192, 216–221 (2015). 10.1016/j.jsb.2015.08.008

123 Galkin, V. E., Orlova, A., Vos, M. R., Schroder, G. F. & Egelman, E. H. Near-atomic resolution for one state of F-actin. Structure 23, 173–182 (2015). 10.1016/j.str.2014.11.006

124 Zivanov, J., Nakane, T. & Scheres, S. H. W. A Bayesian approach to beam-induced motion correction in cryo-EM single-particle analysis. IUCrJ 6, 5–17 (2019). 10.1107/S205225251801463X

125 Scheres, S. H. RELION: implementation of a Bayesian approach to cryo-EM structure determination. J Struct Biol 180, 519–530 (2012). 10.1016/j.jsb.2012.09.006

126 Pettersen, E. F. et al. UCSF ChimeraX: Structure visualization for researchers, educators, and developers. Protein Sci 30, 70–82 (2021). 10.1002/pro.3943

127 Pintilie, G. D., Zhang, J., Goddard, T. D., Chiu, W. & Gossard, D. C. Quantitative analysis of cryo-EM density map segmentation by watershed and scale-space filtering, and fitting of structures by alignment to regions. Journal of Structural Biology 170, 427–438 (2010). 10.1016/j.jsb.2010.03.007

128 Jumper, J. et al. Highly accurate protein structure prediction with AlphaFold. Nature 596, 583–589 (2021). 10.1038/s41586-021-03819-2

129 Meng, E. C., Pettersen, E. F., Couch, G. S., Huang, C. C. & Ferrin, T. E. Tools for integrated sequence-structure analysis with UCSF Chimera. Bmc Bioinformatics 7 (2006). 10.1186/1471-2105-7-339

130 Liebschner, D. et al. Macromolecular structure determination using X-rays, neutrons and electrons: recent developments in Phenix. Acta Crystallogr D Struct Biol 75, 861–877 (2019). 10.1107/S2059798319011471

131 Afonine, P. V. et al. Real-space refinement in PHENIX for cryo-EM and crystallography. Acta Crystallogr D Struct Biol 74, 531–544 (2018). 10.1107/S2059798318006551

132 Emsley, P., Lohkamp, B., Scott, W. G. & Cowtan, K. Features and development of Coot. Acta Crystallogr D Biol Crystallogr 66, 486–501 (2010). 10.1107/S0907444910007493

133 Williams, C. J. et al. MolProbity: More and better reference data for improved all-atom structure validation. Protein Sci 27, 293–315 (2018). 10.1002/pro.3330

134 Weber, M. S. et al. Structural heterogeneity of cellular K5/K14 filaments as revealed by cryo-electron microscopy. Elife 10 (2021). 10.7554/eLife.70307

135 Nickell, S. et al. TOM software toolbox: acquisition and analysis for electron tomography. J Struct Biol 149, 227–234 (2005). 10.1016/j.jsb.2004.10.006

136 Steinert, P. M., Steven, A. C. & Roop, D. R. The Molecular-Biology of Intermediate Filaments. Cell 42, 411–419 (1985). Doi 10.1016/0092-8674(85)90098-4

137 Kocsis, E., Trus, B. L., Steer, C. J., Bisher, M. E. & Steven, A. C. Image Averaging of Flexible Fibrous Macromolecules - the Clathrin Triskelion Has an Elastic Proximal Segment. Journal of Structural Biology 107, 6–14 (1991). Doi 10.1016/1047-8477(91)90025-R

138 Zhang, X. W. Python-based Helix Indexer: A graphical user interface program for finding symmetry of helical assembly through Fourier-Bessel indexing of electron microscopic data. Protein Sci 31, 107–117 (2022). 10.1002/pro.4186

139 Davies, D. B., Saenger, W., Danyluk, S. S., Federation of European Biochemical Societies. & North Atlantic Treaty Organization. Scientific Affairs Division. Structural molecular biology: methods and applications. (Plenum Press, 1982).

140 Martins, B. et al. Unveiling the polarity of actin filaments by cryo-electron tomography. Structure 29, 488–+ (2021). 10.1016/j.str.2020.12.014

141 Landrum, M. J. et al. ClinVar: improving access to variant interpretations and supporting evidence. Nucleic Acids Res 46, D1062–D1067 (2018). 10.1093/nar/gkx1153

142 Sievers, F. et al. Fast, scalable generation of high-quality protein multiple sequence alignments using Clustal Omega. Mol Syst Biol 7, 539 (2011). 10.1038/msb.2011.75

